# Visible light-triggered NO generation from Naphthalimide-based probe for photoreceptor-mediated plant root growth regulation

**DOI:** 10.1101/550004

**Authors:** Suprakash Biswas, Neha Upadhyay, Debojyoti Kar, Sourav Datta, Apurba Lal Koner

## Abstract

An efficient visible light-triggered nitric oxide (NO) releasing fluorescent molecule is designed and synthesized by coupling 2,6-dimethyl nitrobenzene moiety at the *peri*-position of 1, 8-naphthalimide through an alkene bond. The NO-releasing ability is investigated in details using various spectroscopic techniques, and the photoproduct was also characterized. Further, the photo-generated NO has been employed to examine the effect of photoreceptor-mediated NO uptake on plant root growth regulation.

---

Nitrogen monoxide is considered one of the most important signaling agents, exists in three interrelated chemical forms including nitrosonium cation (NO^+^), nitric oxide (NO•), and nitroxyl anion (NO^−^).^1-2^ Nitric oxide (NO) exists as a highly reactive free radical or reactive oxygen species (ROS), sometimes also referred to as reactive nitrogen species (RNS), performs as an important multitasked signaling agent in various biological processes.^3-5^ NO has also been identified as an endothelium-derived relaxing factor (EDRF) in blood vessels and the nitric oxide synthase is mainly responsible for NO production in living systems.^6-10^ Besides, several diseases such as schizophrenia, Alzheimer’s disease, and cancer are related to malfunction of NO signaling and its dynamics.^11-13^ For the past few decades, researchers also found that NO performs a crucial role in plant growth.^14-15^ In the metabolism of inorganic nitrogenous compounds in higher plants and nitrogen-fixing organism, NO is generated as a pivotal intermediate.^16^ Recent literature on plant biology reveals that NO influences primary root development through the initiation of cell-cycle genes and patterns of cellulose synthesis.^17-18^ Additionally, being an essential signaling agent, deficiency of NO affects auxin biosynthesis, transport, and signaling.^19^ The highly reactive nature of NO and its tremendous biological importance fascinated the researchers for *in-situ* generation of NO in biological systems.^20-21^ For potential therapeutic applications in living systems, the NO-donors must release NO in a time-controlled and site-specific manner.^22-23^ To date, numerous NO-donors such as 1-hydroxy-2-oxo-3-(aminoalkyl)-1-triazenes, 4-alkyl-2 hydroxyimino-5-nitro-3-hexenes, *etc.* are reported in the literature.^24-25^ Among them, photo-triggered NO-donors secure extraordinary attention to the researchers due to the release of NO with high spatiotemporal control. Miyata group reported 6-Nitrobenzo[*α*] pyrene derivative that generates NO in the presence of visible light.^26^ Later, they reported a series of NO-releasing molecules, comprising of a hindered nitrobenzene derivative, but all those molecules are poorly water-soluble.^27^ However, the recently reported light-induced NO-donors suffer from ultraviolet and two-photon excitation.^28 29-31^ Therefore, there is an urgent need of a visible-light triggered efficient NO-donor with considerable water-solubility.

In this contribution, we have designed and synthesized a naphthalene monoimide-based fluorescent molecule (Ni-NO_2_) for a visible light-induced NO generation. Thus, we have attached a sterically hindered nitrobenzene moiety, i.e., 2, 6-dimethyl nitrobenzene, at the *peri*-position of 1, 8-naphthalimide ring through an alkene spacer. Subsequently, elaborative spectroscopic investigations were performed to understand the visible light-triggered NO release and further employed to comprehend the effect of NO in photoreceptor-mediated plant root growth regulation. The design strategy of Ni-NO_2_ is based on photo-isomerization of nitro group of the sterically-hindered nitrobenzene moiety, connected to the naphthalimide ring. Considering the photoisomerization possibility, we have synthesized Ni-NO_2_ according to the above synthetic scheme (Scheme 1), the detailed synthetic scheme is shown in Scheme S1. Density functional study reveals that the nitro group is in non co-planar conformation with a twist angle of *ca.* 42°, with respect to the benzene ring due to the steric hindrance of two *ortho*-methyl groups (Scheme S2a-b and Fig. S1). Upon photo-irradiation, the twisted nitro group rearranges and isomerizes to nitrite ester,^28^ owing to the generation of NO (as shown in Scheme S2c).

**Scheme 1.**
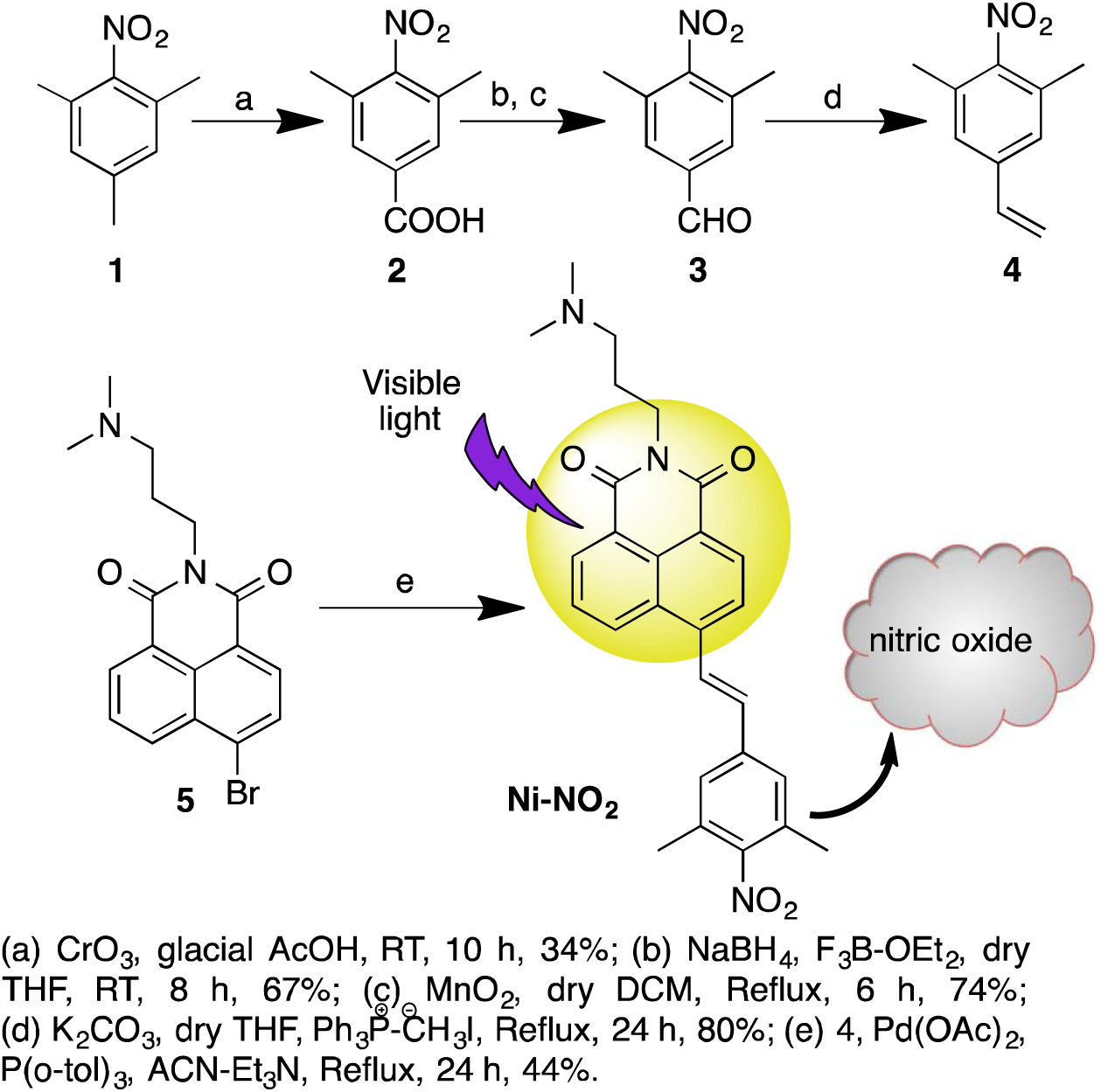
Synthesis of Ni-NO_2_ for visible-light triggered NO generation

After the synthesis and characterization, we validated the optical purity by comparing UV-Vis. and excitation spectra in phosphate buffer (PB, pH 6, Fig. 1a). Ni-NO_2_ has an absorption maximum in PB at around 400 nm, whereas it emits around 517 nm (Fig. 1a) with a large Stokes shift of 117 nm. To understand the swiftness of NO-releasing properties of Ni-NO_2_, we have kinetically monitored its fluorescence properties under visible-light excitation (113 Lux) in 1 mM PB at pH 6. The fluorescence intensity of Ni-NO_2_ gets attenuated upon visible-light mediated excited-state-isomerization of the nitro group. Interestingly, within 20 minutes of irradiation, we have obtained a plateau in the emission intensity indicating completion of NO release (Fig. 1b). The fluorescence quenching is possibly due to the formation of the phenolic group and its interaction with the solvent.^32-33^ Further, to verify the effect of light intensity on the NO release rate, we have performed kinetic monitoring of NO release experiment using a neutral density filter and recorded the emission spectra of Ni-NO_2_. As expected, we obtained a plateau after 30 min (Fig. S2-3) confirming the role of light intensity for NO release. Furthermore, we have performed a modified Griess assay to validate NO generation through the photolysis of Ni-NO_2_.^27^ The photo-induced production of NO was verified by the detection of NO_2_^−^, arising from the auto-oxidation of NO in PB medium.^34^ The appearance of red color solution having absorption at 542 nm supports the diazo-coupling reaction of NO_2_^−^ with the Griess reagents (GR, Scheme S3). To perform this assay, we have taken 50-150 *μ*M of Ni-NO_2_ in CHCl_3_ and excited by 400 nm light for ∼50 min, and then the mixture was allowed to react with GR. Finally, the absorption spectrum of the red color solution was measured. The increase optical density at 542 nm upon increasing concentration of Ni-NO_2_, signifies the successful generation of NO (Fig. 2a). Further, to testify the photo-triggered generation of NO, an EPR experiment was also performed using 2-phenyl-4, 4, 5, 5,-tetramethylimidazoline-1-oxyl-3-oxide (PTIO) as a spin trap for NO.^29,35^ After the photo-irradiation of a mixture of PTIO and Ni-NO_2_for 30 min the EPR spectrum was recorded.

**Fig. 1.**
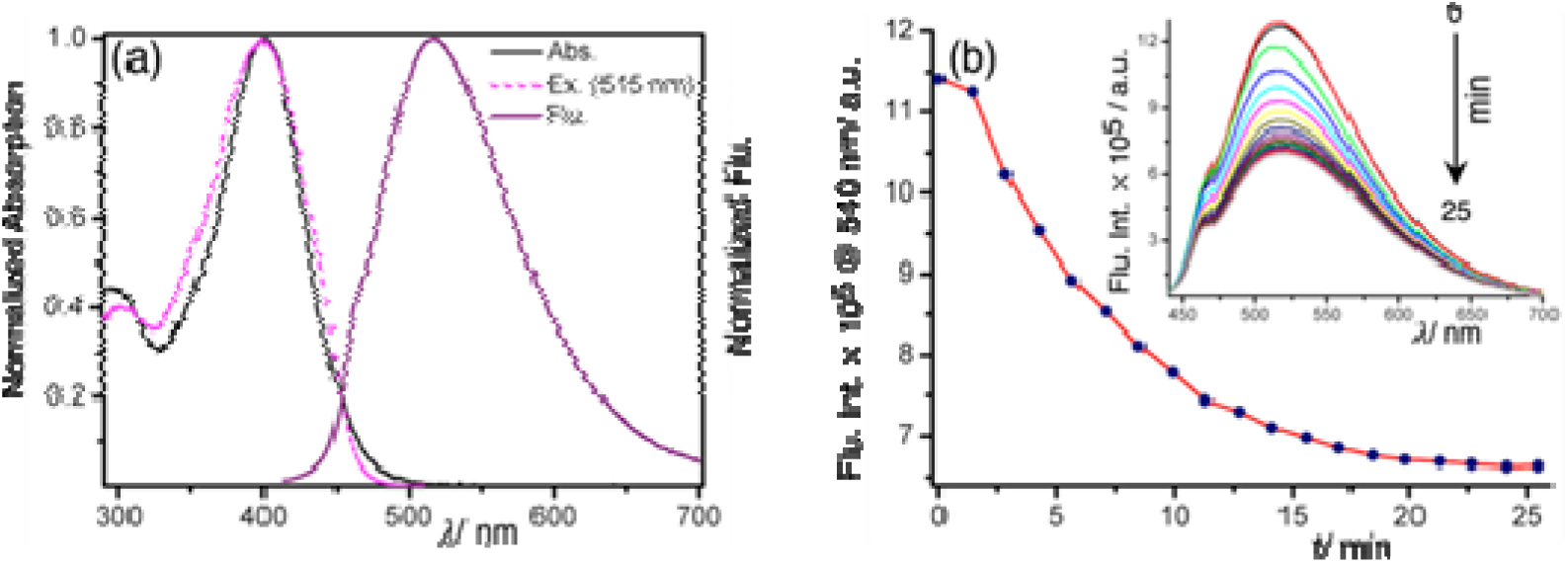
(a) The absorption, excitation, and emission spectra of Ni-NO_2_ in PB at pH 6.0, (b) kinetic monitoring of fluorescence intensity of 1 *μ*M Ni-NO_2_ at 540 nm in PB upon excitation of 410 nm light; inset shows fluorescence spectra of the same at different time interval

**Fig. 2.**
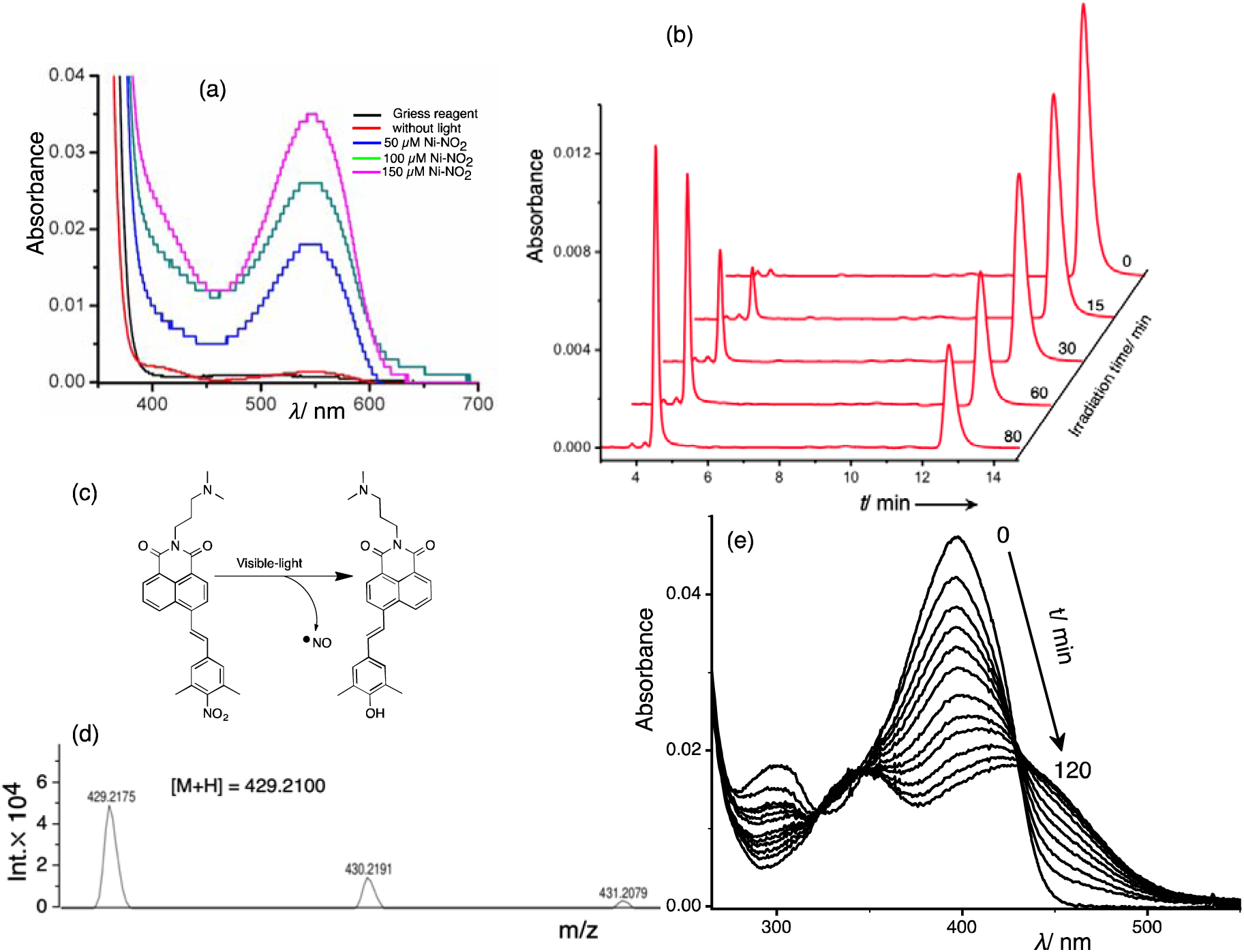
(a) Absorption spectra of the reaction between Griess solution with a photo-irradiated Ni-NO_2_ solution, (b) chemical structure of the photo-generated product of Ni-NO_2_ and (c) the mass-spectrum of the product after photodecomposition, (d) HPLC profile of photo-triggered NO generation experiments and (e) time-dependent absorption spectra of Ni-NO_2_ (2 *μ*M) in DMSO upon photo-irradiation using white light.

The characteristic EPR spectrum further confirms the generation of NO from Ni-NO_2_ (Fig. S4). Additionally, we have performed a HPLC experiment to monitor the progress of the reaction. For HPLC measurement, we have irradiated 10 *μ*M of Ni-NO_2_ in acetonitrile using 390 nm light at 0, 15, 30, 60, and 80 min interval (see Fig. 2b and SI for details). A linear decrease in the absorption intensity corresponding to the reactant having an elution time 12.8 minute and subsequent increase in the absorption intensity for the product at an elution time of 4.6 minutes were observed (Fig. 2b). The photodecomposition quantum yield of NO release was measured as *ca.* 0.036 in acetonitrile (for details see S1, Fig. S5-7). The photo-degraded product was also identified by mass spectrometry (Fig. 2c-d). Further, the photo-reaction was also monitored by absorption spectroscopy in different time interval in DMSO (Fig. 2e) and in other solvents (see Fig. S8). The absorption spectra in DMSO reveal a decrease in the absorption maxima at 400 nm with the generation of a new absorption peak at around 427 nm which also confirms the formation of photo-generated product.^33^

After the spectroscopic characterization of the NO release in the presence of visible-light, we have investigated the effect of NO on plant roots and root-hair growth. A thorough literature survey suggests that NO inhibits root growth in a dose-dependent manner.^36^ Hence, to determine the effect of NO released from Ni-NO_2_, we used *Arabidopsis thaliana* as a model plant and its root growth inhibition as a functional assay. Arabidopsis seeds (Col-0 ecotype) were inoculated on growth medium supplied with 0, 0.2,1.0, 5.0,10.0, 30.0 *μ*M of Ni-NO_2_. The NO-donor Ni-NO_2_ was added inside the upper cover of plates at desired concentrations (Fig. S9). Seedlings were grown on Ni-NO_2_ free agar medium in the lower portion of the plate to avoid side effects of NO-donor reagents.^37^ One set of plates was kept under cycling white light (110 *μ*M m^−2^ sec^−1^) while the other set was kept under darkness. After six days seedlings grown under light and containing Ni-NO_2_ in the upper lid showed significantly reduced primary root length as compared to the plate without Ni-NO_2_ (Fig. 3a). On the other hand, seedlings grown in the dark did not exhibit any significant difference in root length in the presence or absence of Ni-NO_2_ (Fig. 3a). These results suggest that the photo-activation of Ni-NO_2_ promotes the release of NO that causes root growth inhibition in plants. In order to evaluate the efficiency of Ni-NO_2_ derived NO on root growth inhibition, we used sodium nitroprusside (SNP), a well-established NO-donor, and compared the effects on primary root growth inhibition (Fig. 3b-c).^38^ At similar concentrations of these two NO-donors the root elongation defect was either comparable or more severe in Ni-NO_2_ compared to SNP (Fig. 3b-c). Further, we examined the effect of Ni-NO_2_ and SNP with varying fluence levels of cycling white light, 35, 85 and 150 *μ*M m^−2^ sec^−1^ (Fig. 3b and Fig. S10a-b). At all fluence levels, both the NO-donors showed reduced primary root length in a dose-dependent manner. We also determined the effect of continuous white light and flash of white light (1 h at 150 *μ*M m^−2^ sec^−1^) on NO-donors by using different doses of Ni-NO_2_ and SNP (0, 2 and 20 *μ*M) and exposing them to continuous or flash of white light. Both the light conditions exhibited an inhibitory effect on primary root growth (Fig. 3c and Fig. S11a, S12a). Further, we investigated the effect of monochromatic blue light on NO-donors by exposing plants to continuous blue light (150 *μ*Mm^−2^ sec^−1^) and a flash of blue light (1 h at 150 *μ*M m^−2^ sec^−1^, see S1). Both Ni-NO_2_ and SNP markedly inhibited primary root length under continuous blue light (SI, Fig. S11b, S12b). Under flash of blue light, Ni-NO_2_ at 2 *μ*M concentration promoted primary root length while at 20 *μM* concentration it showed inhibitory effect on root growth (Fig. S12b, S13b). This might be due to cell elongation effect at a low concentration of NO under blue light.^39^ We also examined the effect of different concentrations (0-10 *μ*M) of Ni-NO_2_ and SNP on root-hair growth when exposed to continuous white light (110 *μ*M m^−2^ sec^−1^). Root hair growth is inhibited (reduced root hair length and number) in a dose-dependent manner (Fig. 4a-c). This result suggests that the regulatory effect of Ni-NO_2_ on root growth of Arabidopsis seedlings is light dependent. Upon exposure to light, the photolysis of Ni-NO_2_ releases NO which inhibits root growth. Light-mediated release of NO also activates various photoreceptors in plants. Phytochrome (PHY) and cryptochrome (CRY), red and blue light receptors respectively are reported to be NO-responsive. NO release causes nuclear accumulation of *phyB-9* which positively regulates primary root growth inhibition.^40^ To verify, if the NO-induced primary root growth inhibition is photoreceptor mediated, we quantified root length in the photoreceptor mutants *phyB-9* and *cry1cry2* (Fig. 5 and S14-18). Our results indicate that *phyB-9* and *cry1cry2* exhibit hyposensitivity to the NO released by 2 *μ*M Ni-NO_2_. At 10 *μ*M concentration of Ni-NO_2_ the root growth inhibition was also observed in the mutants especially under white light (WL) possibly due to the activation of other photoreceptors in the single mutant (Fig. 5a-b). Under the monochromatic blue light condition, *cry1cry2* double mutantdisplay almost insensitivity at a higher concentration indicating that NO-induced primary root growth inhibition is photoreceptor mediated (Fig. 5c-d).

**Fig. 3.**
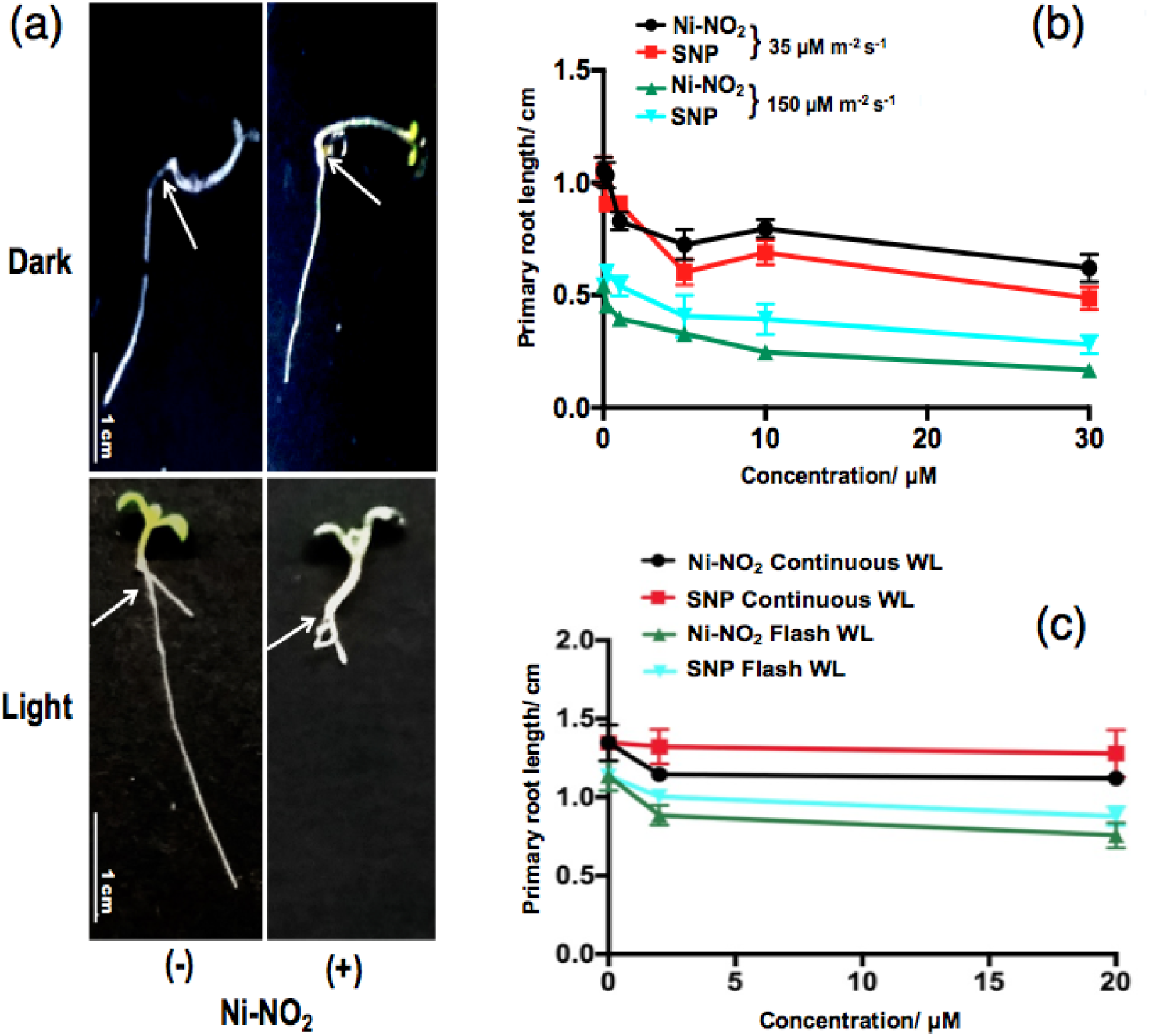
Response of Arabidopsis (Col-0) roots to Ni-NO_2_ under dark and light, (a) 6 days old Arabidopsis seedlings grown on MS media plates supplemented with 0,10*μ*M Ni-NO_2_ in continuous dark and cycling white light at 110 *μ*M m^−2^ s^−1^ fluence level. Part of seedling below white arrow shown in (a) indicates root that is inhibited by Ni-NO_2_ in light due to the release of NO. (b) Primary root length of Col-0 seedlings treated with 0, 0.2,1, 5, 10, 30 *μ*M Ni-NO_2_ and SNP at two different fluence levels 35 and 150 *μ*M m^−2^ s^−1^ of cycling white light (c) and in continuous white light (110 *μM* m^−2^ s^−1^) and flash of white light (110 *μ*M m^−2^ s^_1^ for 1 hour). Scale bar, 1 cm in (a).

**Fig. 4.**
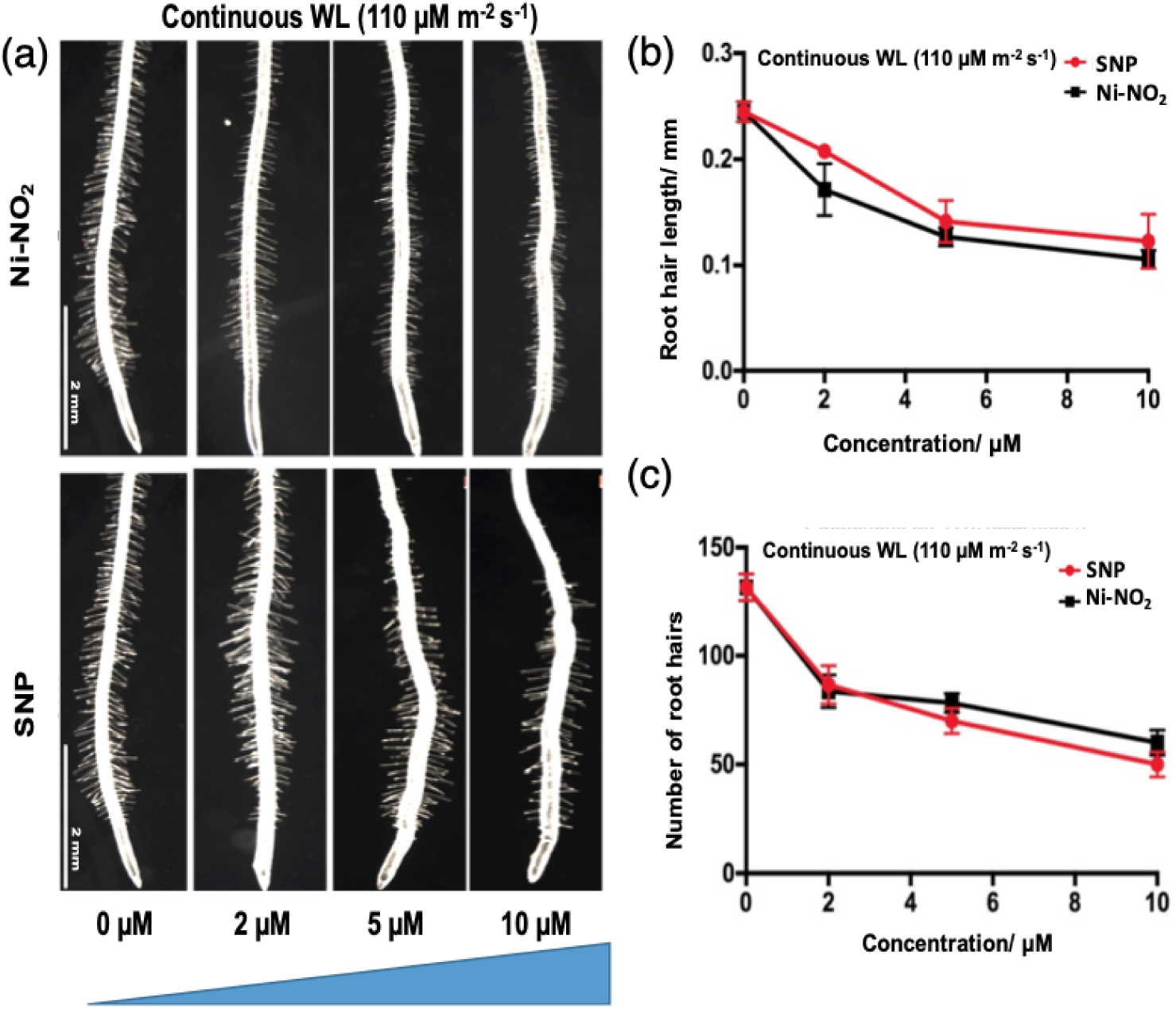
Inhibition of root hair development by Ni-NO_2_ under white light, (a) Root hair image of 6 days old Col-0 seedlings grown on medium supplemented with 0, 2, 5, 10 *μM* Ni-NO_2_ and SNP in continuous white light at 110 *μ*M m^−2^ s^−1^ fluence level (b) Root hair length measurement of Col-0 seedlings treated as described in (a), (c) Root hair number measurement of Col-0 seedlings treated as described in (a). Mean values and S.E. were calculated from at least 30 seedlings. Scale bar, 2 mm in (a).

**Fig. 5.**
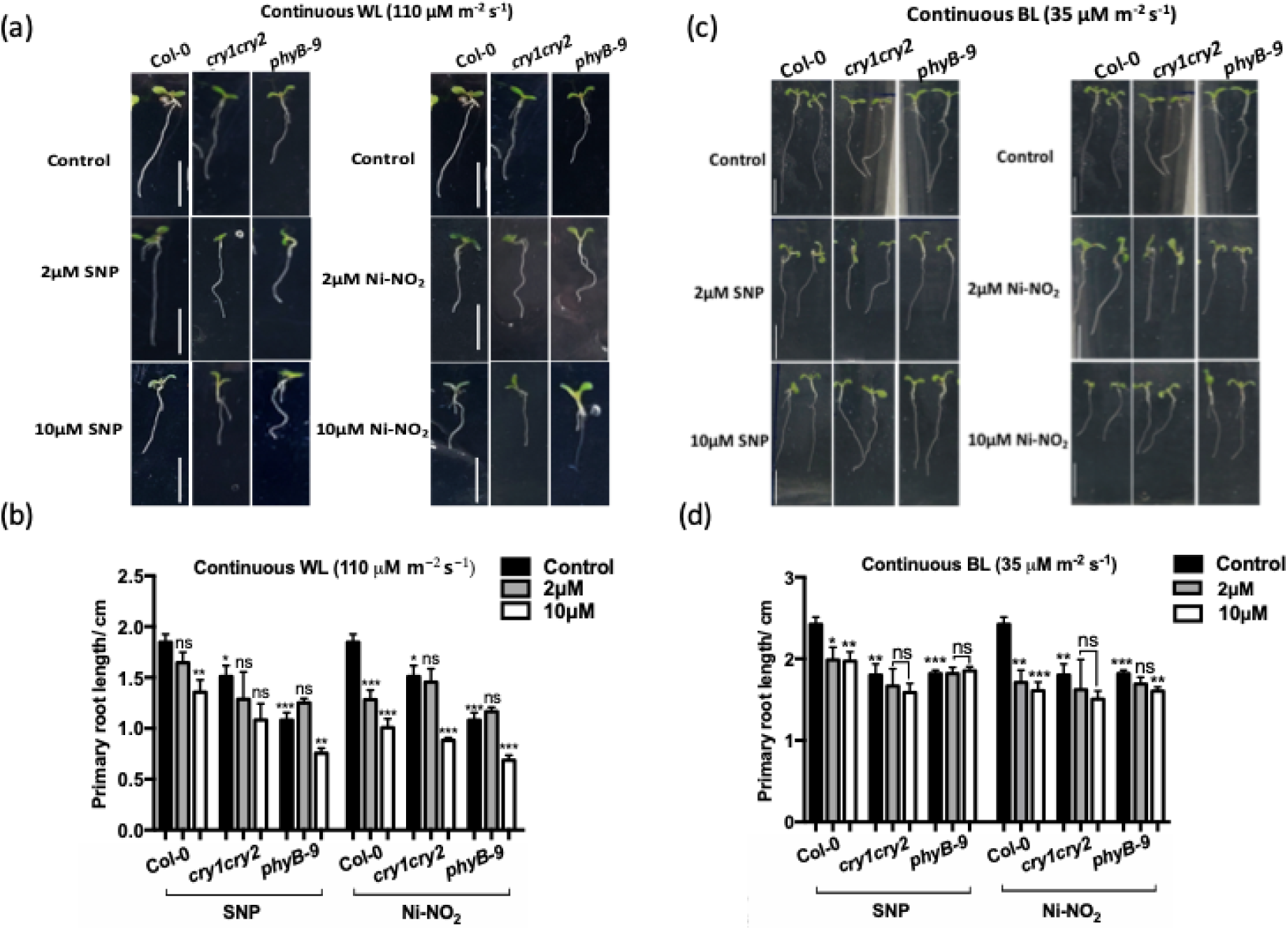
Response of WT, *cry1cry2* and *phyB-9* mutants to NO under continuous white and blue light (a) WT, *cry1cry2* and *phyB-9* seedlings growing on medium supplemented with SNP and Ni-NO_2_ at various concentration under continuous WL at 110 *μM* m^−2^ s^1^ fluence. (b) Root length of WT, *cry1cry2* and *phyB-9* seedlings treated as in (a), (c) WT, *cry1cry2* and *phyB-9* seedlings growing on medium supplemented with SNP and Ni-NO_2_ at 0, 2 and 10 *μ*M concentration under continuous blue light at 35 *μ*M m^−2^ s^1^ fluence. (d) Root length of WT, *cry1cry2* and *phyB-9* seedlings treated as in (c). Mean value and SE were calculated from at least 20 seedlings. Significant difference was analyzed student’s t-test with Welch’s corrections compared to WT under same condition are indicated by asterisks: *P <.05, ** *P* <.01, *** *P* <.001.

In conclusion, we have designed and developed a naphthalene monoimide-based fluorescent NO-donor Ni-NO_2_ by attaching 2,6 dimethyl nitrobenzene at the *peri*-position of 1,8-naphthalimide ring through an alkene bond. The synthesized molecule releases NO upon photoexcitation with visible light and the NO generation was confirmed by Griess assay, EPR spin trapping, HRMS and optical spectroscopic investigation in various solvents. Further, the progress of the reaction was monitored using HPLC. The photodecomposition quantum yield of Ni-NO_2_ in acetonitrile was also determined. The fluorescence intensity of the probe gets quenched by releasing NO in aqueous solution and its get saturated within 20 min signifying the rapid release of NO. Finally, the released NO was employed for Arabidopsis root growth assay either by continuous or flash of white and blue light irradiation. Our results indicate NO-mediated regulation of primary root and root hair growth by Ni-NO_2_ in a photoreceptor mediated manner. These findings can potentially open up prospects for generation of an effective weedicide in the future.

## Supporting Information

Materials and methods, synthesis, photodecomposition QY measurement, kinetic study, DFT study, Griess assay, root growth experiments, characterization by UV-Vis., NMR, EPR and mass spectrometry.

## ACKNOWLEDGMENT

We acknowledge the financial support from IISER Bhopal (INGRANT/CHM/2012-017). SB thanks UGC, India for his fellowship. SD would like to thank DBT, India (Ramalingaswami Fellowship, IYBA) and SERB (EMR/2016/000181), India for funding. NU and DK acknowledge MHRD and DBT (BT/PR19193/BPA/118/195/2016), India for fellowship. Authors acknowledge Mr. Anil Raj Narooka for his help with HPLC measurements.

