## supplemental file for "Visible light-triggered NO generation from Naphthalimide-based probe for photoreceptor-mediated plant root growth regulation"

**Table of Contents**

1. Materials and instruments………..............................................................................................................3

2. Theoretical calculation…………...............................................................................................................3

3. Plantmaterials and growth conditions…....................................................................................................4

4. Root length measurement…………...........................................................................................................4

5. Scheme1: Synthetic scheme………….......................................................................................................4

6. Synthetic procedure …………...............................................................................................................5-7

7. DFT optimized structure and plausible mechanism of NO release........................................................7-8

8. Figure S1: Energy optimized Kuhn-Sham HOMO and LUMO of Ni-NO_2_….........................................8

9. Z-matrix for the optimized configuration of Ni-NO_2_……...................................................................9-10

10. Figure S2: Time-dependent emission spectra of Ni-NO_2_ upon photo-excitation with 400 nm light….10

11. Figure S3: Plot of emission intensities of Ni-NO_2_ at 540 nm with different time upon photo irradia...11

12. Figure S4: EPR spectrum of PTIO and the reaction mixture (PTIO with Ni-NO_2_)………...................11

13. Photo-decomposition quantum yield measurement…………………...................................................12

14. Preparation of potassium ferrioxalate trihydrate K_3_[Fe(C_2_O_4_)_3_].3H_2_O……………….........................12

15. Calculation of the Photon flux…………………………………...........................................................13

16. Figure S5: UV-Vis absorption of [Fe(Phen)_3_]^2+^ complex at different time interval…………..............13

17. Figure S6: Plot of number of moles of Fe^2+^ formed against different time of light irradiation….........14

18. Photo-decomposition quantum yield calculation………………….......................................................15

19. Figure S7: Plot of number of moles of product against photon flux………………..............................15

20. HPLC experiment details………………………..............................................................................15-16

43. Figure S29: 1H NMR spectrum of compound 5……............................................................................27

44. Figure S30: 13C NMR spectrum of compound 5……..........................................................................27

45. Figure S31: 1H NMR spectrum of compound NI-NO_2_…….................................................................28

50. Figure S36: APCI HRMS mass spectrum of compound **3b**……...........................................................29

51. Figure S37: APCI HRMS mass spectrum of compound **4**……….........................................................29

52. Figure S38: APCI HRMS mass spectrum of compound **5**…….............................................................30

53. Figure S39: APCI HRMS mass spectrum of compound **NI-NO_2_**……..................................................30

54. References……….……………........................................................................................................30-31

**Experimental Procedures**

**Materials and instruments**:

All the reactions were performed in oven-dried glassware under a nitrogen atmosphere where required and stirred with Teflon-coated magnetic stirring bars. Tetrahydrofuran was distilled over sodium/benzophenone ketyl. All other solvents and reagents were used as received unless otherwise stated. Reaction temperatures above room temperature (298 K) refer to oil bath temperature. Thin layer chromatography was performed using Merck Silica gel 60 F-254 pre-coated plates and visualized using UV irradiation (λ=254/365 nm). Silica gel from Merck (particle size 100-200 mesh) was used for column chromatography. 1H and 13C NMR spectra were recorded on Bruker 400 MHz spectrometers with operating frequencies of 100 MHz for 13C. Chemical shifts (δ) are reported in ppm relative to the residual solvent signal (δ = 7.26 for 1H NMR and δ = 77.0 for 13C NMR). High-resolution mass spectrometry (HRMS) data were recorded on MicrOTOF-Q-II mass spectrometer using chloroform as the solvent. All the reagents and solvents were brought from sigma used without further purification. Water used was of Milli-Q grade using a Millipore water purification system with resistivity 18.2 MΩ, for the phosphate buffer preparation. All absorption spectra and fluorescence measurements were carried out using SHIMADZU spectrophotometer and HORIBA JobinYvon fluorimeter (fluorolog) using 1 cm path length quartz cuvettes. EPR spectra measurement was carried out in BRUKER A-300, X-band EPR spectrometer with modulation amplitude of 3 G, modulation frequency 10 KHz, and central field 3440 G. HPLC experiments were performed using a UV detector from Waters 2489 fitted with a reverse-phase C18 column 5 m, 4.6 mm x 250 mm. Samples were injected using a auto sampler from Waters (part no: 2707). NMR spectroscopic characterization was taken on a BRUKER Advance 400 MHz spectrometer. A 2.0 mM stock solution of Ni-NO_2_ was prepared in DMSO and then further diluted to 1.0 M in phosphate buffer of pH 6.0 to check the absorption and steady-state fluorescence spectra of Ni-NO_2_.

**Theoretical calculation**

Density functional theory calculations were performed on Ni-NO_2_ using Gaussian 09 suite of quantum chemical programs.^1^ Ground-state geometry optimizations were performed with Becke three-parameter exchange functional in conjunction with Lee-Yang-Parr correlation functional (B3LYP)^2-4^ using 6-311G as a basis set.

**Plant Materials and Growth Conditions:**

Arabidopsis thaliana Columbia-0 (Col-0) ecotype, *phyB-9*, *cry1cry2* seeds were used in this study. Seeds were surface-sterilized with 5% (v/v) bleach solution for about 3 min, rinsed with sterile water at least four times, and then sown on half-strength Murashige and Skoog (1/2 MS) medium containing 0.8% (w/v) agar. After 3 d at 4°C in the dark to synchronize germination, the plates were transferred to a growth chamber with continuous white light (about 110 μM m^–2^ s^–^1) and maintained at 22 °C for 5 days. To avoid possible effects of chemicals other than NO gas on root growth, NO treatments were performed by photochemical release of NO gas from the NO donor Ni-NO_2_ and sodium nitroprusside (SNP), which was mixed with a small amount of growth medium before solidification and added inside the upper cover of the plates at the desired concentrations. Seedlings grew on Ni-NO_2_ and SNP-free agar medium in the lower portion of the plate. In this way, we could ensure that the effect on the seedlings was due to NO rather than to Ni-NO_2_ and SNP.

**Root length measurement:**

Seedlings were laid horizontally on the agar plates, digital pictures were taken, and primary root length and root hair length of at least 30 seedlings were measured using ImageJ software.

**Synthetic Scheme:**

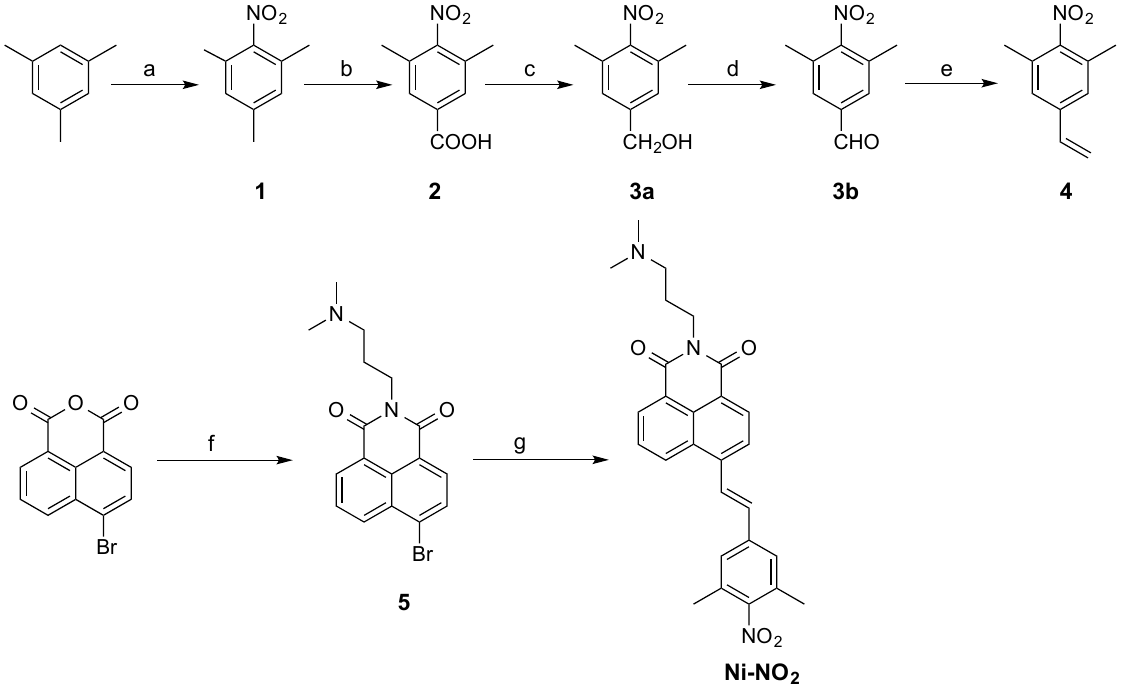

**Scheme S1:** Synthesis pathway of **Ni-NO_2_**

Reagent and conditions: (a) HNO_3_, AcOH, Acetic anhydride, 0-5 °C, then 50 °C for 30 min; (b) CrO_3_, glacial AcOH, RT, 10 h; (c) NaBH_4_, BF_3_-Et_2_O, dry THF, RT, 8 h; (d) MnO_2_, dry DCM, Reflux, 6 h; (e) K_2_CO_3_, dry THF, [Ph_3_P-CH_3_]I, Reflux, 24 h; (f) 3-(Dimethylamino)-1-propylamine, EtOH, Reflux, 2 h; (g) 6, Pd(OAc)_2_, P(*o*-tol)_3_, ACN-Et_3_N, 24 h, Reflux.

**Synthesis procedure:**

1,3,5-trimethyl-2-nitrobenzene (**1**): 10 g (86 mmol) of 1,3,5-Trimethylbenzene and 14.5 g (141.5 mmol) of acetic anhydride were poured in a round bottle flask and cooled to 0 °C with an ice bath. A mixture of 5.6 mL of fuming nitric acid (SG: 1.52) and 5.5 g (91.5 mmol) of glacial acetic acid was added dropwise to the mesitylene-acetic anhydride solution by keeping the temperature at 0 °C. After the addition was completed, the flask was removed from ice bath and allowed to stir at room temperature for two hours. After that, the reaction mixture was warmed to 50 °C for 30 minutes. After completion of the reaction, total reaction mixture was poured into ice-cold water and then saturated NaHCO_3_ solution was added to the mixture to adjust the pH over 7.0. After that the organic layer was extracted with ethyl acetate and dried over MgSO_4_. The crude reaction product was subject to column chromatography with 1: 10 ethyl acetate/ hexane mixture and finally we got a solid (10.25 g) yellowish crystalline product (yield 74.6%). ^1^H NMR (400 MHz, CDCl_3_) δ 6.91 (s, 2H), 2.31 (s, 3H), 2.27 (s, 6H). ^13^C{1H} NMR (100 MHz, CDCl_3_) δ 149.90, 140.41, 129.54, 21.00, 17.50. HRMS (APCI) m/z [M+H^+^] calculated for C_9_H_11_NO_2_ 165.0790 Da and found 165.0784 Da

3,5-dimethyl-4-nitrobenzoic acid (**2**): 12 g (120 mmol) CrO_3_ was first dissolved 140 mL glacial acetic acid and stirring vigorously to make a suspension. In a separate round bottom flask, 6 g (36.35 mmol) of compound **1** was dissolved in 15 mL of glacial acetic acid and the solution was added dropwise to the suspension of CrO_3_ under vigorous stirring. After complete addition, the whole reaction mixture was allowed to stir at room temperature for 15 h. After completion of the reaction whole reaction mass was transferred to ice-cold water and a white precipitate appeared. The precipitate was collected by filtration and washed several times with water and the precipitate was dissolved in CHCl_3_. The CHCl_3_ layer was washed with 2 N NaOH, and the aqueous layer was acidified with conc. HCl. After acidification a white precipitate appeared which was extracted with CHCl_3_. The CHCl_3_ layer was washed with brine and dried over MgSO_4_, and finally the pure product was collected by solvent evaporation which affords 2.4 g of white solid product (yield 34.5%). ^1^H NMR (400 MHz, CDCl_3_) δ 7.90 (s, 2H), 2.38 (s, 6H) ^13^C{1H} NMR (100 MHz, CDCl_3_) δ 170.30, 155.05, 130.79, 130.25, 129.99, 17.26. HRMS (APCI) m/z [M-H] calculated for C_9_H_9_NO_4_ 194.0532 Da and found 194.0448 Da

(3,5-dimethyl-4-nitrophenyl)methanol (**3a**): To a slurry of NaBH_4_ ( 875 mg, 23 mmol ) in dry THF ( 15 mL), compound **2** (1.5 g, 7.7 mmol) in 10 mL THF was added drop wise, followed by the addition of 0.98 mL of F_3_B-Et_2_O ( 875 mg, 6.15 mmol) by maintaining inert atmosphere with N_2_. The reaction mixture was allowed to stir at room temperature for 10 h. After completion of the reaction ice-cold water was added to the reaction mixture slowly and the crude product was extracted with CHCl_3_. The CHCl_3_ layer was washed with brine and dried over Na_2_SO_4_. The crude product was subject to column chromatography in 20 % ethyl acetate and hexane which gave 1.1 g of solid yellow product (yield 79%). ^1^H NMR (400 MHz, CDCl_3_) δ 7.12 (s, 2H), 4.68 (d, 2H, J = 5.44 Hz), 2.31 (s, 6H), 1.83 (t, 1H, J = 5.78 Hz) ^13^C{1H} NMR (100 MHz, CDCl_3_) δ 150.94, 142.73, 130.00, 126.71, 64.05, 17.36. HRMS (APCI) m/z [M+] calculated for C_9_H_11_NO_3_ 181.0739 Da and found 181.0733 Da

3,5-dimethyl-4-nitrobenzaldehyde (**3b**): To a slurry of MnO_2_ (4.09 g, 47 mmol) in DCM, compound **3a** (850 mg, 4.7 mmol) was added and the reaction was allowed to reflux for 6 h, by maintaining N_2_ atmosphere. After completion of the reaction, the reaction mixture was filtered through celite and the crude product was subject to column chromatography in 20% ethyl acetate and hexane mixture. The column chromatography gives 621 mg light yellowish solid product (yield 74 %). ^1^H NMR (400 MHz, CDCl3) δ 9.99 (s, 1H), 7.66 (s, 1H), 2.38 (s, 6H), ^13^C{1H} NMR (100 MHz, CDCl_3_) δ 190.71, 155.06, 136.62, 130.61, 130.09, 17.16, HRMS (APCI) m/z [M+H^+^] calculated for C_9_H_9_NO_3_ 179.0582 Da and found 180.0655 Da

1,3-dimethyl-2-nitro-5-vinylbenzene (**4**): To a suspension of compound **3b** (400 mg, 2.2 mmol) and K_2_CO_3_ (397 mg, 2.9 mmol) in anhydrous THF (7 mL) [Ph_3_PCH_3_]I was added (1.16 g, 2.8 mmol) and stirred under reflux for 24 hours. After that, the reaction mass was concentrated under reduced pressure and the residue was dissolved in water. The aqueous layer was extracted with ethyl acetate. The ethyl acetate layer was dried with MgSO_4_, filtered and the crude product subject to silica gel column chromatography with 20 % ethyl acetate and hexane mixture. The column chromatography afforded pure product as a yellow liquid (yield 80 %). ^1^H NMR (400 MHz, CDCl_3_) δ 7.16 (s, 2H), 6.67 (dd, 1H, J = 10.90 Hz, 17.55 Hz), 5.81 (d, 1H, J = 17.57 Hz), 5.38 (d, 1H, J = 10.87 Hz), 2.34 (s, 6H), ^13^C{1H} NMR (100 MHz, CDCl_3_) δ 150.89, 139.27, 135.21, 129.99, 126.58, 116.56, 17.50, HRMS (APCI) m/z [M+] calculated for C_10_H_11_NO_2_ 177.0790 Da and found 177.0863 Da

6-bromo-2-(3-(dimethylamino)propyl)-1H-benzo[de]isoquinoline-1,3(2H)-dione (**5**): To a solution of 4-Bromo-1,8-naphthalic anhydride (500 mg, 1.8 mmol) in EtOH 0.24 mL (1.9 mmol) of 3-(Dimethylamino)-1-propylamine was added and the resulting mixture was refluxed for 2 h and monitored by TLC. After completion of the reaction, the reaction mass was cooled to room temperature and concentrated under vacuum this affords solid white product. Further, the solid crude product was purified by silica gel column chromatography with 20% EtOAc and Hexane mixture, provides 565 mg of solid white product (yield 83.4%). ^1^H NMR (400 MHz, CDCl_3_) δ 8.66 (d, 1H, J = 7.28 Hz), 8.58 (d, 1H, J = 7.31 Hz), 8.42 (d, 1H, J = 8.32 Hz), 8.05 (d, 1H, J = 7.84 Hz), 7.85 (t, 1H, 7.84 Hz), 4.23 (t, 2H, J= 6.88 Hz), 2.43 (t, 2H, J= 6.64 Hz), 2.24 (s, 6H), 1.91 (m, 2H) ^13^C{1H} NMR (100 MHz, CDCl_3_) δ 163.52, 133.23, 132.00, 131.19, 131.08, 130.63, 130.21, 129.01, 128.07, 123.13, 122.27, 57.24, 45.37, 38.95, 26.03, HRMS (APCI) m/z [M+H^+^] calculated for C_17_H_17_BrN_2_O_2_ 360.0473 Da and found 361.0536 Da

(E)-6-(3,5-dimethyl-4-nitrostyryl)-2-(3-(dimethylamino)propyl)-1H-benzo[de]isoquinoline-1,3(2H)-dione (**Ni-NO_2_**): To a high pressure bottle compound 5 (77 mg , 0.21 mmol), palladium (II) acetate (2.4 mg, 5 mol%) and tri-o-tolyl phosphine ( 6.51 mg, 10 mol%) were combined and 2.5 mL of the solvent pair (triethylamine 0.75 mL / acetonitrile 1.75 mL) were added. To the former mixture a solution of compound 4 (50 mg, 0.27 mmol) in 0.5 mL acetonitrile was added, the bottle was sealed after bubbling with nitrogen for 10 min. The system was heated to 105 °C, for refluxing for 24 h. After completion of the reaction whole reaction mass was concentrated under vacuum and to the reaction mixture water was added and the organic product was extracted with CHCl_3_. After Drying with MgSO_4_, a yellow solid crude product was isolated and finally the product was purified by silica gel column chromatography with 2% MeOH and CHCl_3_ which affords 44.0 mg of yellow product (yield 44%). ^1^H NMR (400 MHz, CDCl_3_) δ 8.65 (d, 1H, J = 7.28 Hz), 8.60 (d, 1H, J = 7.14 Hz), 8.56 (d, 1H, J = 8.58 Hz), 7.98 (d, 1H, J = 7.75 Hz), 7.91 (d, 1H, J = 16.07 Hz), 7.81 (dd, 1H, J = 8.51 Hz, 7.26 Hz), 7.37 (s, 2H), 7.24 (d, 1H), 4.24 (t, 2H, J = 6.88 Hz), 2.48 (t, 2H, J = 7.36 Hz), 2.39 (s, 6H), 2.29 (s, 6H), 1.95 (m, 2H) ^13^C{1H} NMR (100 MHz, CDCl_3_) δ 164.17, 163.94, 151.48, 140.60, 138.21, 133.35, 131.34, 131.00, 130.54, 129.81, 129.63, 128.69, 127.42, 127.00, 126.17, 124.32, 123.23, 122.12, 57.22, 54.30, 38.83, 26.01, 17.74 HRMS (APCI) m/z [M+H^+^] calculated for C_27_H_27_N_3_O_4_ 457.2002 Da and found 458.2074 Da

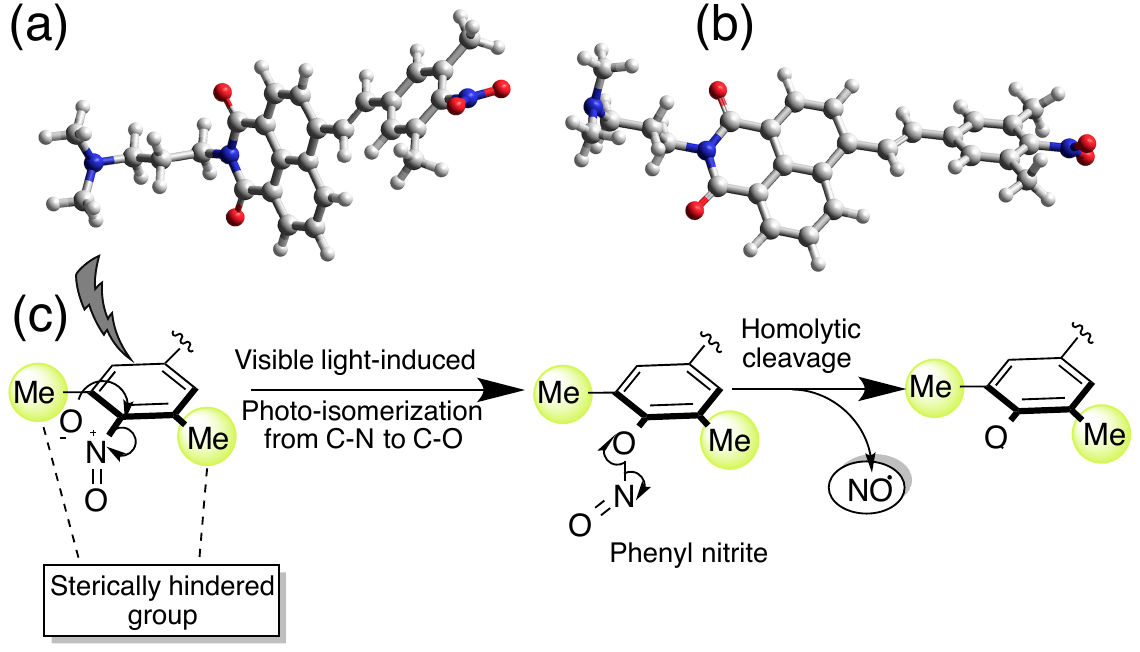

**Scheme S2**: (a-b) DFT optimized structure at the B3LYP/6-311G level shows the twisting of the nitro group due to the steric hindrance of *ortho*-methyl groups, (c) schematic representation of the plausible mechanism of NO release from Ni-NO_2_, under visible light exposure.

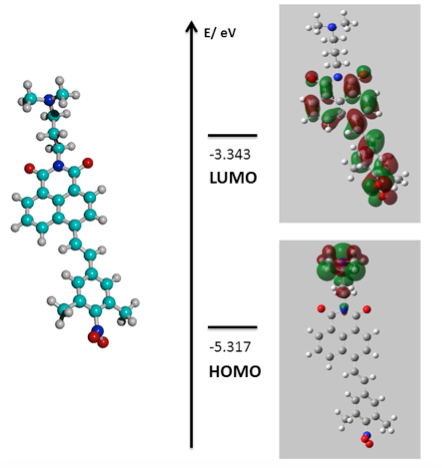

**Figure. S1**: Energy optimized Kuhn-Sham HOMO and LUMO of Ni-NO_2_

Symbolic Z-matrix for the optimized configuration of Ni-NO2: Charge = 0 Multiplicity = 1

C	1.20802	1.26617	-2.20239

C	2.14789	1.78673	-0.0121

C	1.13305	2.78522	0.15276

C	0.16833	2.99748	-0.86663

C	0.21174	2.23712	-2.02354

H	1.24525	0.72523	-3.13724

C	1.08805	3.5604	1.34354

H	-0.52619	2.42227	-2.79075

C	2.01282	3.34989	2.3517

C	3.00075	2.35717	2.20664

H	1.95172	3.95547	3.24419

H	3.71	2.1897	3.00517

C	-0.87782	4.02533	-0.7106

C	0.0502	4.59735	1.5215

O	-0.0135	5.30919	2.54675

O	-1.73544	4.24696	-1.5928

C	3.06474	1.59262	1.05758

H	3.81961	0.82402	0.98263

C	2.17746	1.01556	-1.23101

C	3.22457	0.01263	-1.45896

H	4.1756	0.19721	-0.97474

C	3.08459	-1.10177	-2.2078

H	2.11083	-1.32383	-2.63212

C	4.11786	-2.10346	-2.48886

C	3.73958	-3.31313	-3.09532

C	5.4777	-1.91465	-2.1846

C	4.65361	-4.33226	-3.37487

H	2.69972	-3.47106	-3.35043

C	6.44449	-2.88099	-2.46275

H	5.8059	-0.98817	-1.73493

C	-1.92259	5.80751	0.6514

H	-2.79714	5.46742	0.10261

H	-2.15103	5.8561	1.71331

N	-0.88147	4.76254	0.48491

C	-1.46679	7.1806	0.13754

H	-0.5636	7.4798	0.67347

H	-1.21707	7.09982	-0.92269

C	-2.57307	8.244	0.3376

H	-3.48004	7.92088	-0.18259

H	-2.82343	8.30476	1.40119

C	-2.15855	9.77725	-1.57078

H	-1.23571	9.36289	-2.01204

H	-3.00967	9.29886	-2.05724

H	-2.19011	10.8417	-1.81143

C	-1.23307	10.31164	0.64181

H	-1.26622	11.37465	0.39541

H	-1.43463	10.20881	1.70891

H	-0.20536	9.9569	0.45261

N	-2.26123	9.60391	-0.12161

C	6.00387	-4.08775	-3.05141

N	6.9918	-5.12548	-3.34289

O	8.11003	-4.77619	-3.83065

O	6.68969	-6.33303	-3.09633

C	7.88771	-2.5953	-2.11455

H	8.37346	-3.44263	-1.63085

H	7.94265	-1.74073	-1.441

H	8.47087	-2.3734	-3.00744

C	4.15683	-5.60998	-4.01125

H	4.1183	-6.42046	-3.28421

H	4.80312	-5.95085	-4.81983

H	3.15665	-5.45668	-4.41492

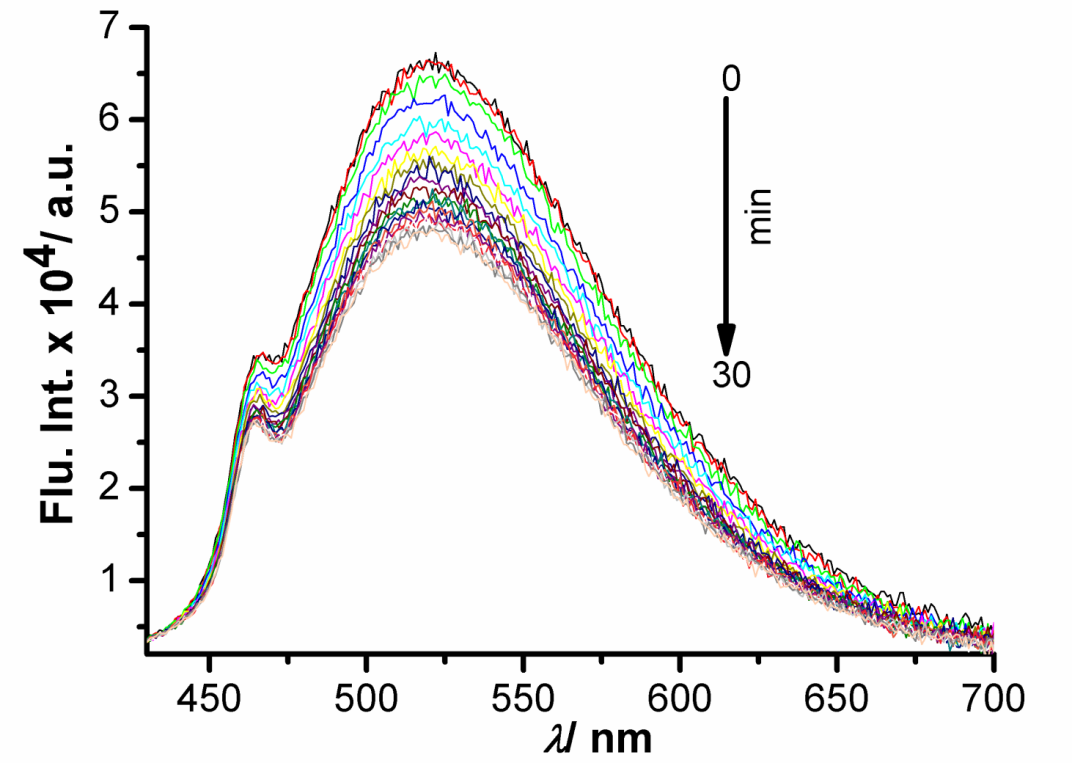

**Figure S2.** Emission spectra of Ni-NO_2_ (1.0 *μ*M) at different time, when the probe is photo-excited with 400 nm of light in phosphate buffer of pH 6.0, through a neutral density filter, which reduces 10 times the incident light intensity.

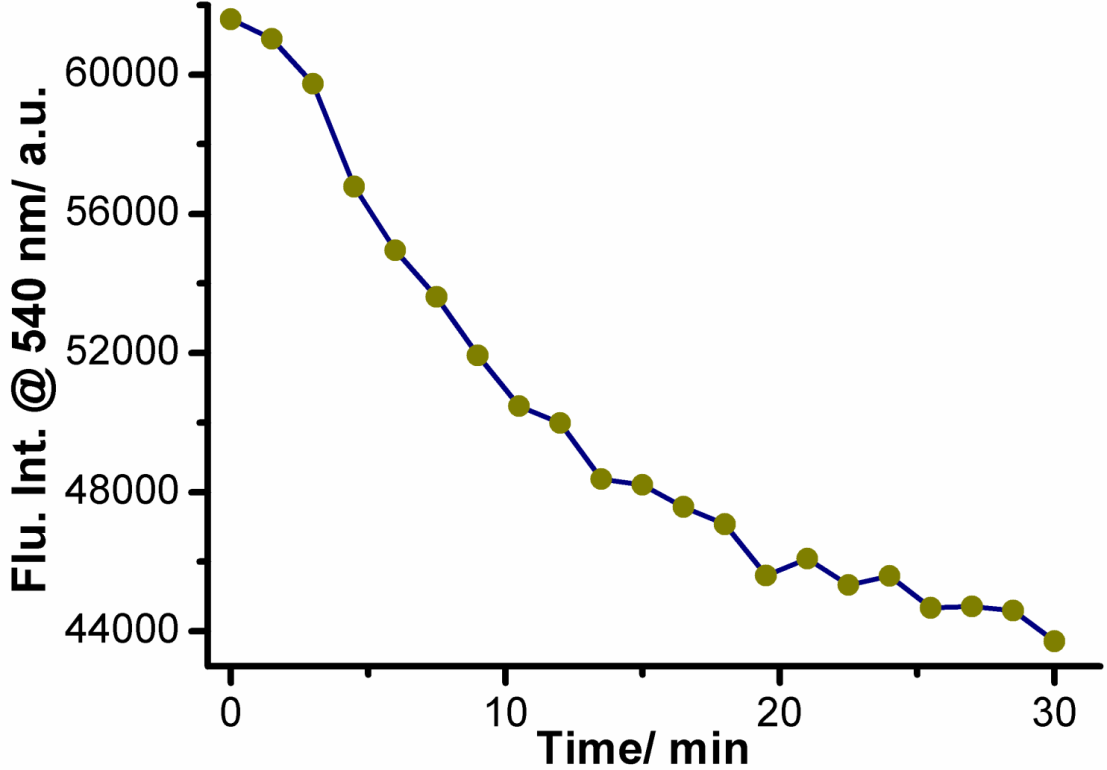

**Figure S3.** Plot of emission intensities at 540 nm measured in different time when 1.0 *μ*M of Ni-NO_2_ is photo-irradiated for 30 min with 400 nm of light through neutral density filter which reduces 10 times the incident light intensity in phosphate buffer of pH 6.0.

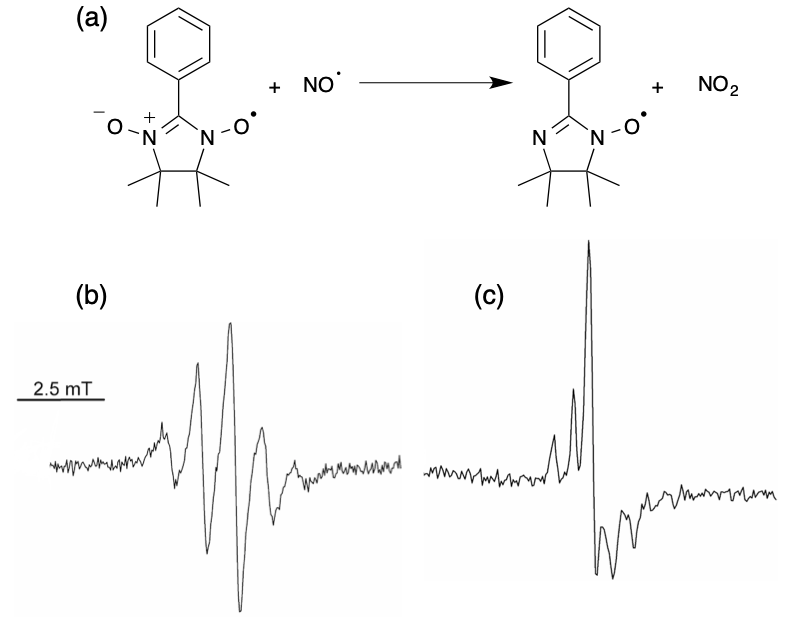

**Figure S4.** (a) Chemical reaction between PTIO and NO. EPR spectrum of 5 *μ*M PTIO in DMSO (b) without Ni-NO_2_ and (c) with photo-irradiated solution of 10 *μ*M of Ni-NO_2_. Photo-decomposition quantum yield measurement:

**Photo-decomposition quantum yield measurement:**

We have measured the photo-decomposition quantum yield of Ni-NO_2_ using potassium ferrioxalate trihydrate K_3_[Fe(C_2_O_4_)_3_].3H_2_O salt as a standard by following the modified literature reported procedure.^5^ To perform this we have prepared an aqueous solution of ferrioxalate actinometer was prepared and kept in dark. In presence of light, the potassium ferrioxalate solution photo-decomposes and transformed to ferrous oxalate anions from ferric oxalate anions. Subsequently, these ferrous oxalate anions can react with 1,10-phenanthroline to form the [Fe(Phen)_3_]^2+^ complex. The concentration of this [Fe(Phen)_3_]^2+^ complex can be determined by monitoring the UV-Vis. absorbance having a maxima at 510 nm in Milli-Q water. The number of photons absorbed by the ferrioxalate actinometer solution is equivalent to the no. of moles of [Fe(Phen)_3_]^2+^ complex formed. Henceforth, we can determine the photon flux which is directly related to our photo-induced NO release and hence the quantum yield of the photo-decomposition reaction can be determined. Detailed procedure of the quantum yield measurement is as follows. All the reactions were carried out at room temperature (298 K).

**Preparation of K_3_[Fe(C_2_O_4_)_3_].3H_2_O salt**^6^

Oxalic acid dihydrate (3.78 g) was taken in a conical flask (carefully wrapped with aluminium foil) and 10 mL of distilled water was added to make the solution of oxalic acid. Next, 3.36 g of KOH was added to it slowly with vigorous stirring. Afterwards, 1.62 g of FeCl_3_ was added to the above solution under constant stirring condition. The reaction was complete within 5-10 minutes as evident from the appearence of dark green color. Later, the solution was kept in dark for 1 h to obtain green crystals of K_3_[Fe(C_2_O_4_)_3_].3H_2_O. The crystals were filtered, washed with cold distilled water, dried and stored in dark.

**Determination of amount of photon absorbed by potassium ferrioxalate solution**:

1 mL of potassium ferrioxalate trihydrate (K_3_[Fe(C_2_O_4_)_3_].3H_2_O) solution (25 mM, prepared in 1N H_2_SO_4_) and 1 mL Ni-NO_2_ (10 *μ*M, prepared in Acetonitrile) were taken in two different 1 mL quartz cuvette having path length 1 cm. Both the cuvettes were exposed seperately one after another using 390 nm monochromatic light. A 450 W Hg lamp was used as light source for *λ*= 390 nm light. Further, the actinometer solution was irradiated for 0, 100, 200 and 300 second intervals respectively. The reaction mixture was also irradiated for 15, 30, 60 and 80 min intervals respectively and the no. of moles of product formed in each case were measured by HPLC.

**Photon flux calculation:**

After each interval 0.1 mL of the irradiated potassium ferrioxalate trihydrate solution was transferred into a 1 mL volumetric flask containing 0.2 mL of 1,10-phenanthroline solution (5.84 mM, prepared in 60 mM sodium acetate buffer having H_2_SO_4_ concentration 2.7 N). Then the volumetric flask volume was adjusted with distilled water, shaken and kept in dark for 1h for complete complexation. After that, 1 mL of the complex was taken in 1 mL of quartz cuvette and the absorption spectra of the solutions were reorded. The absorption at *λ* = 510 nm corresponds to the amount of amount of Fe^2+^ formed in the photodecomposition process (Figure S5).

A blank solution was also prepared with 0.2 mL of 1,10-phenanthroline solution, 0.1 mL of actinometer (without any irradiation) and the volume was adjusted with distilled water in a 1 mL of volumetric flask. The absorbance spectrum of the blank solution was also measured (see inset of Figure S5).

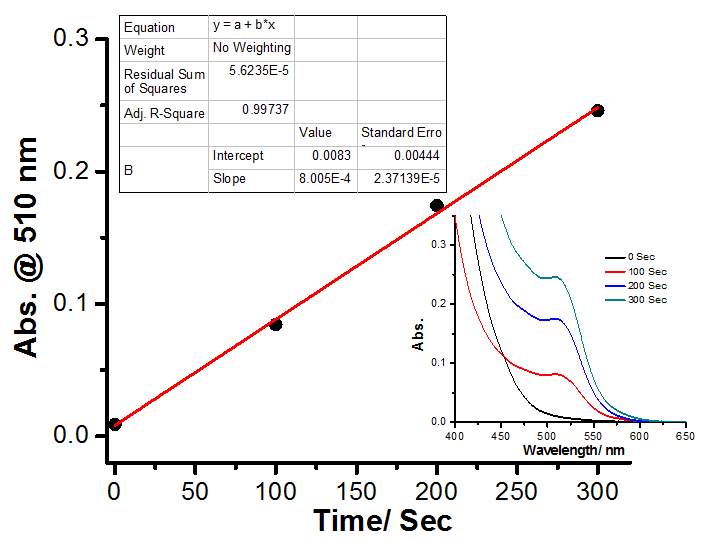

**Figure S5.** Plot of UV-Vis absorption of [Fe(Phen)^3^]^2+^ complex at different time interval of irradiation and the inset is showing the corresponding absorption spectra. According to Beer’s law, the number of moles of Fe^2+^ formed is determined using the following formula:

$${Fe}^{2+}=\frac{V_{1}V_{3}\Delta A( 510 nm)}{{10}^{3}V_{2}l\varepsilon( 510 nm)}$$

Where: V1 = Irradiated volume of actinometer solution (1 mL).

V2 = The aliquot taken from the irradiated actinometer solution for the estimation of Fe^2+^ ions (0.1 mL). V3 = The volume of the solution after complexation with 1,10-phenanthroline (1 mL).

ε = Molar extinction coefficient of [Fe(Phen)_3_] ^2+^ complex (11100 Lmol-1cm-1) at 510 nm.^7^

l = Optical path-length of the quartz cuvette (1 cm).

ΔA (510 nm) = Difference of absorbance between the irradiated actinometer solution and the blank actinometer solution stored in dark.

No. of moles of Fe^2+^ formed was plotted against time (t) and fitted linearly. The slope (dx/dt) of the line is equal to the number of moles of Fe^2+^ formed per unit time (Figure S6). Hence, we are getting the slope is 0.0725 × 10^-8^ moles/ sec (Figure S6).

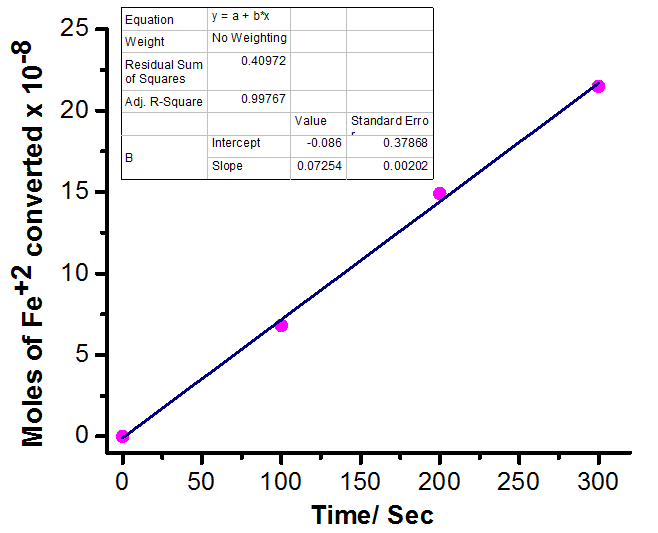

**Figure S6.** Plot of number of moles of Fe^2+^ formed against different time of light irradiation

Afterwards, the no. of incident photons per unit time (F) can be measured by the following equation

$$\phi\left( \lambda\right)= \frac{\frac{dx}{dt}}{F(1-{10}^{-A(\lambda)})}$$

Where, dx/dt = The number of moles of Fe^2+^ formed per unit time is 0.0725×10^−8^ moles/ sec.

Φ(λ) = The quantum yield for the conversion of Fe^3+^ to Fe^2+^ at 390 nm is 1.1.^8^

A(λ) = Absorbance of the ferrioxalate actinometer stock solution at a wavelength of 390 nm.

The obtained value of the absorbance is 0.964.

Hence, the numbers of the photones (F) per unit time, was calculated from the above equation is 7.394× 10^-10^ moles/ sec.

**Determination of photo-decomposition quantum yield (Φ) of NI-NO_2_ by the photoinduced nitric oxide release:**

The no. of moles of product formed at 15, 30, 60 and 80 min are 0.022×10^−6^, 0.046×10^−6^, 0.096×10^−6^ and 0.126×10^−6^ respectively (determined form HPLC chromatogram, by calculating the area under the curve in the chromatogram, Figure 2d). Now, the no. of moles of product is plotted against the no. Moles of incident photons from the light source at the 15, 30, 60 and 80 min respectively (Figure 2b) and fitted linearly. The slope of the plot is equal to the quantum yield (Φ) of our reaction, which is 0.0359 (Figure S7)

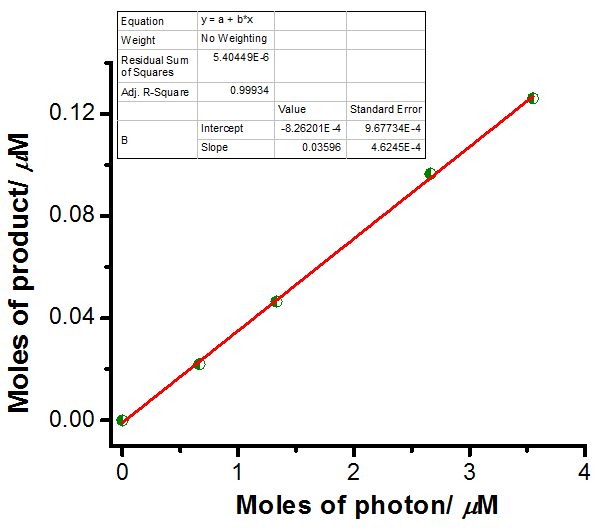

**Figure S7.** Plot of number of moles of product formed against photon flux.

**HPLC experiment:**

This HPLC experiment was performed to calculate the photodecomposition quantum yield as well as the anount of product converted from Ni-NO_2_ upon exposure to the visible light. Firstly, 1 mL solution of Ni-NO2 (10 *μ*M) in acetonitrile was taken in a quatz cuvette and irradiated for 0, 15, 30, 60 and 80 min interval respectively with 390 nm of light. After that, the 30 *μ*L of the reaction mixture at the same time interval was taken and subject to inject connected with a reverse phase C18 HPLC column (length 25 mm) using 3:1 ACN and 10 mM phosphate buffer (pH 6.0 ) as mobile phase with a flow rate of 1 mL/ min. The elutant was filtered with 0.22 micron syringe filter before use. The UV detector in HPLC detector was used to monitored the amount of reactant consumed and the amount of product formed by taking the absorption at 365 nm. Here, we found that at t=0 min i.e, without irradiation HPLC chromatogram shows a single peak centered at 12.8 min and the total area under the curve gives the amount of reactant initially. Afterwards, with the increase in the time of irradiation a new peak genetared at 4.6 min and the intensity as wel as the total area under the curve at 4.6 min is increasing with increasing light irradiation time, correspons to the formation of product. Again the absorption at 12.8 min corresponds to the reactant is also going down with the increase in light exposer time (as shown in Figure 2b). This HPLC chromatograms clearly indicats a linear relation for the formation of product from the reactant and the total area under the peaks at 4.6 min provide the amount of product formed at each time of light irradiation.

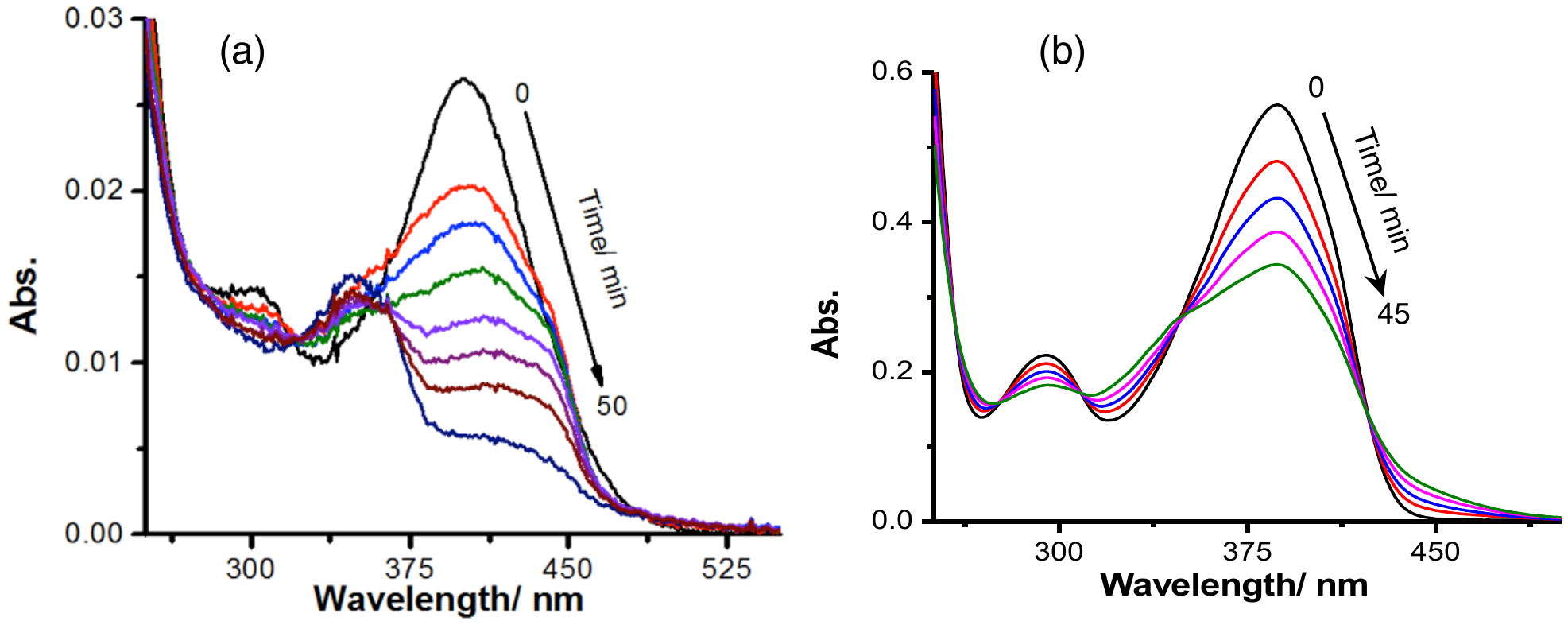

**Figure S8**. Absorption spectra of (a) 2 *μ*M of Ni-NO_2_ in PB at pH 6.0 (b) 20 *μ*M of Ni-NO_2_ in acetonitrile with different irradiation time

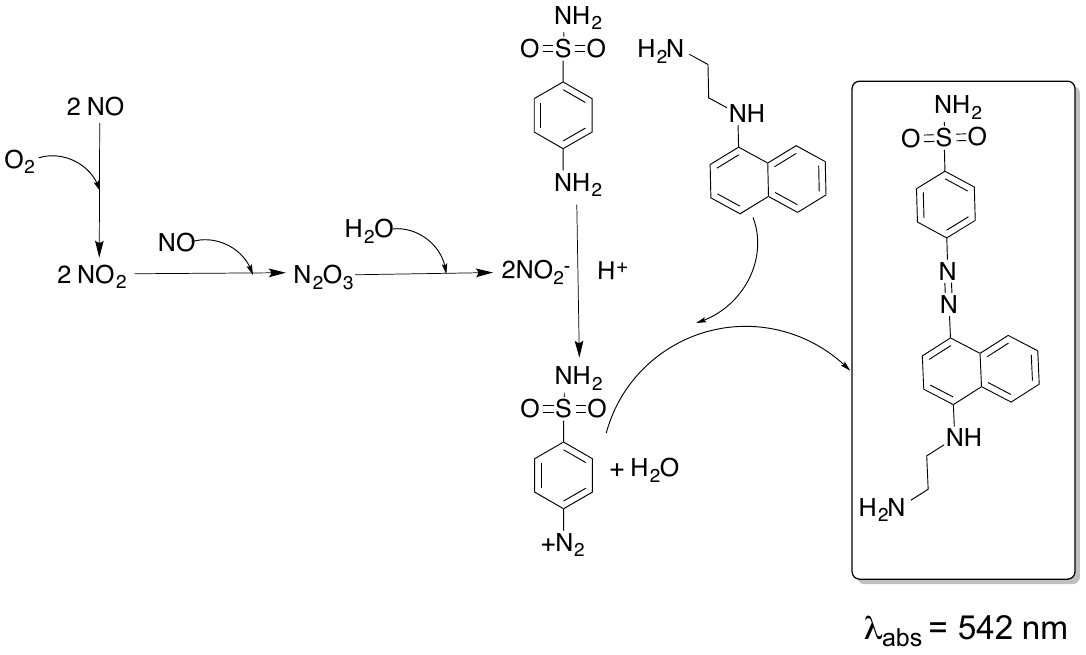

**Scheme S3**. Chemistry associated with Griess assay

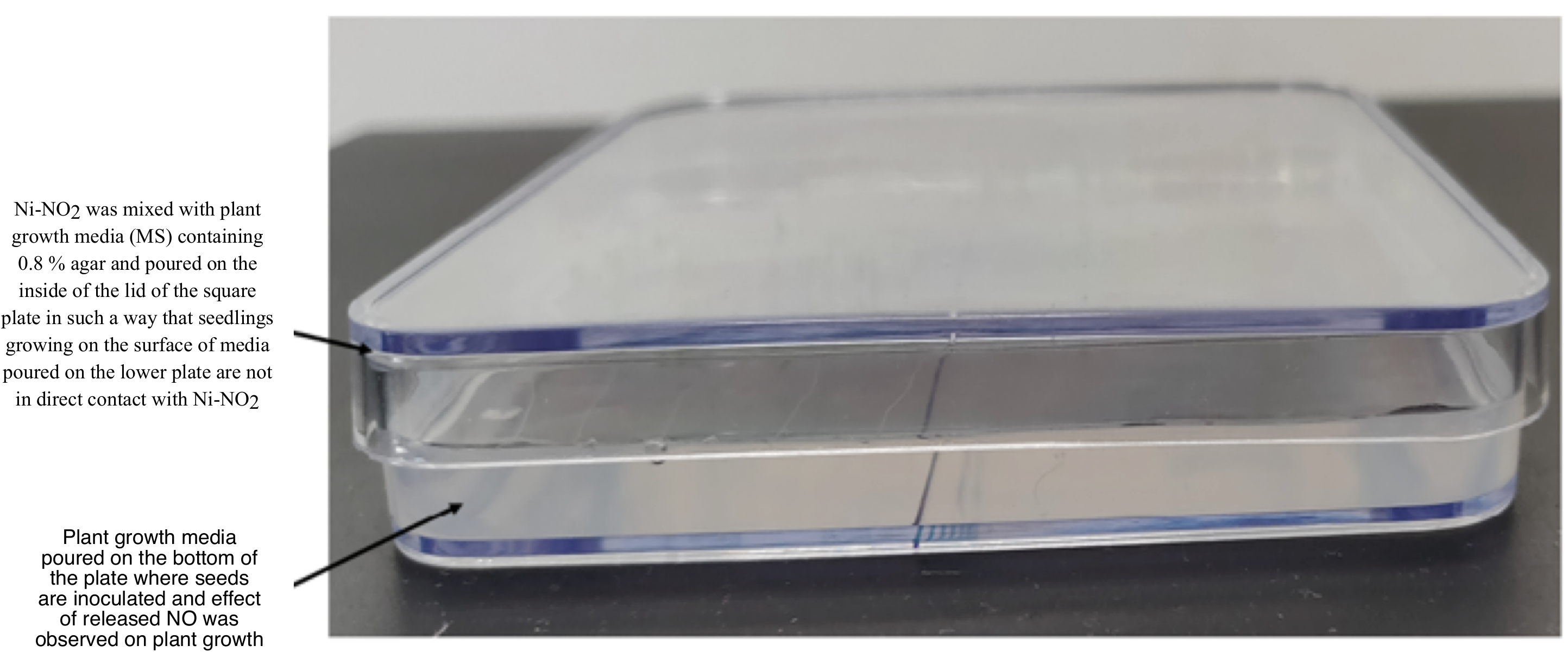

**Figure S9**. Experimental setup for plant growth

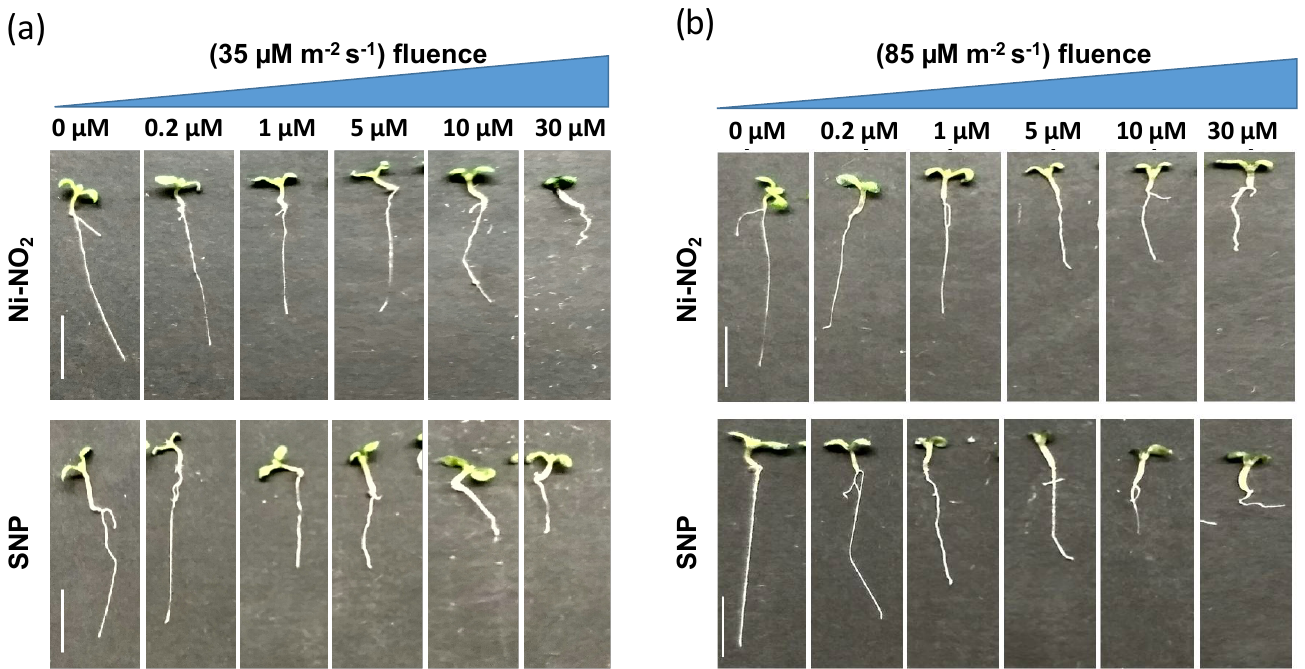

**Figure S10.** Arabidopsis seedlings grown in the presence of NO donors Ni-NO_2_ and SNP under cycling light 6 days old image of Col-0 on growth medium supplemented with 0, 0.2, 1, 5, 10, 30 *µ*M Ni-NO_2_ and SNP at (a) 35 *µ*M m^-2^ s^-1^ and 1 (b) 85 *µ*M m^-2^ s^-1^ of cycling white light. Scale bar, 1 cm (a, b).

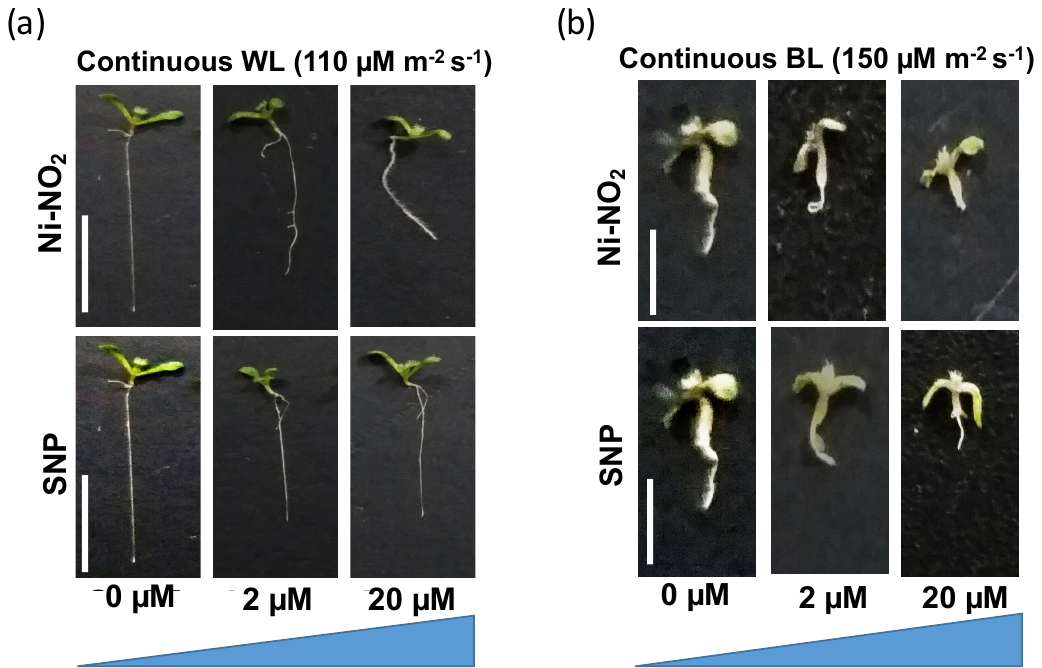

**Figure S11**. Arabidopsis seedlings grown in the presence of NO donors Ni-NO_2_ and SNP under continuous light 6 days old image of Col-0 grown on medium supplemented with 0, 2, 20 *µ*M Ni-NO_2_ and SNP at (a) 110 *µ*M m^-2^ s^-1^ continuous white light (b) 150 *µ*M m^-2^ s^-1^ continuous blue light. Scale bar, 1 cm (a, b).

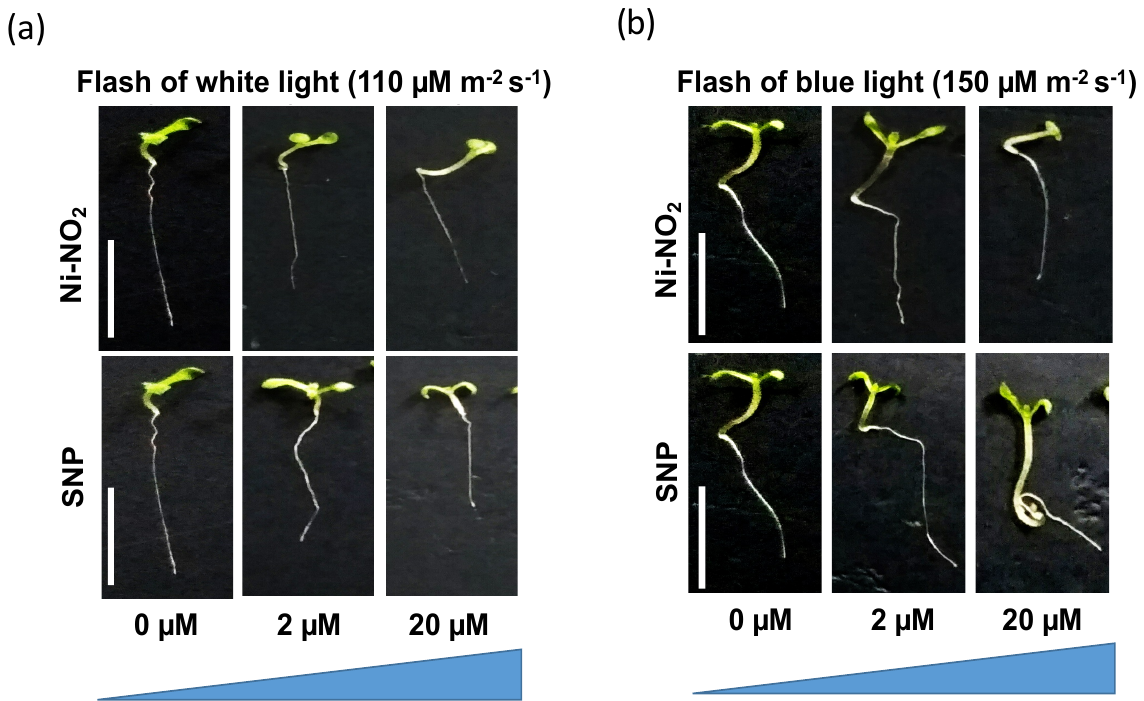

**Figure S12.** Arabidopsis seedlings grown in the presence of NO donors Ni-NO_2_ and SNP under monochromatic light. Flash of white light for 1 hr at 110 *µ*M m^-2^ s^-1^ fluence (a) and flash of blue light for 1 hr at 150 *µ*M m^-2^ s^-1^ fluence (b). Scale bar, 1 cm (a, b).

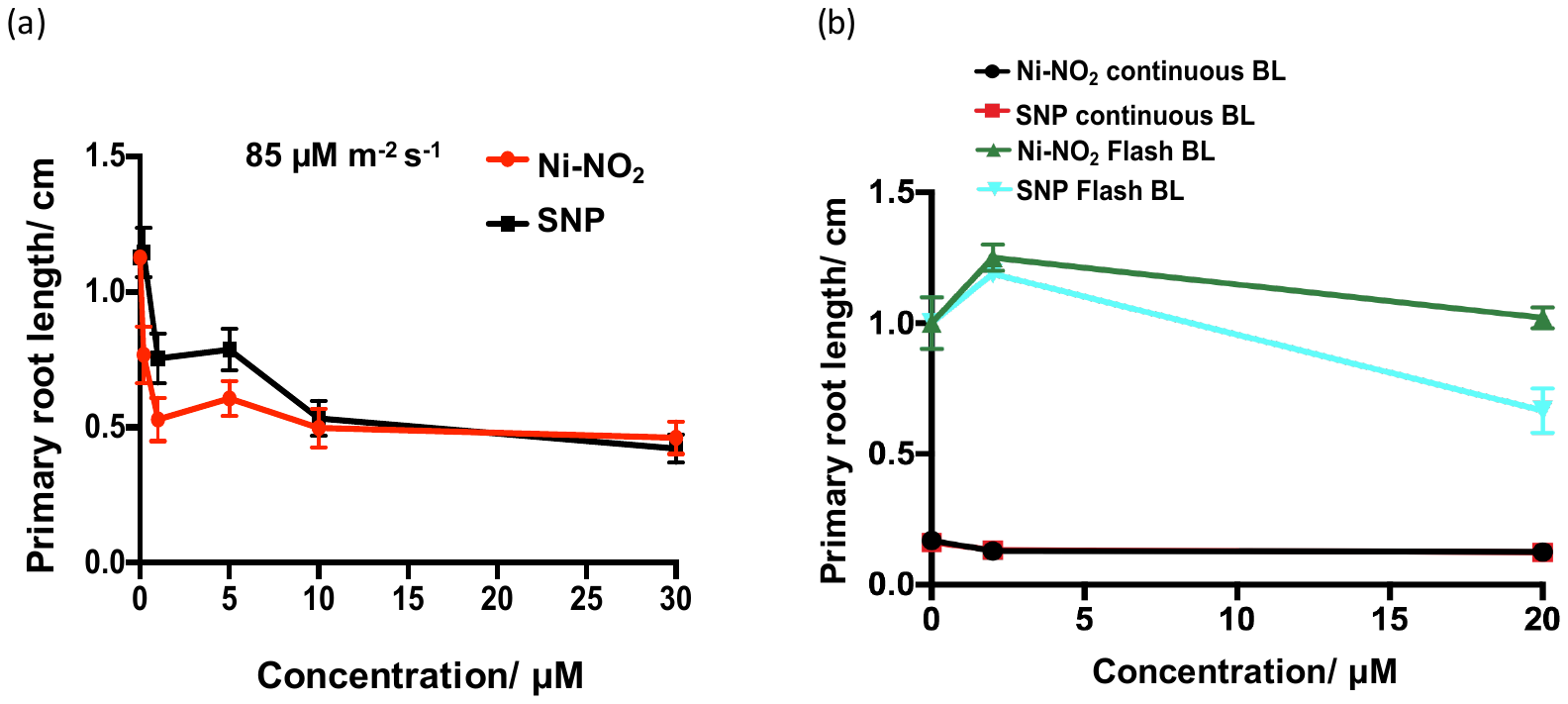

**Figure S13.** (a) Primary root length measurement of Col-0 seedlings treated with 0, 0.2, 1, 5, 10, 30 *µ*M Ni-NO_2_ and SNP in cycling white light at 85 *µ*M m^-2^ s^-1^ fluence; (b) Primary root length measurement of Col-0 seedlings treated with 0, 2, 20 *µ*M Ni-NO_2_ and SNP in continuous and flash of blue light. Mean values and S.E. were calculated from at least 30 seedlings.

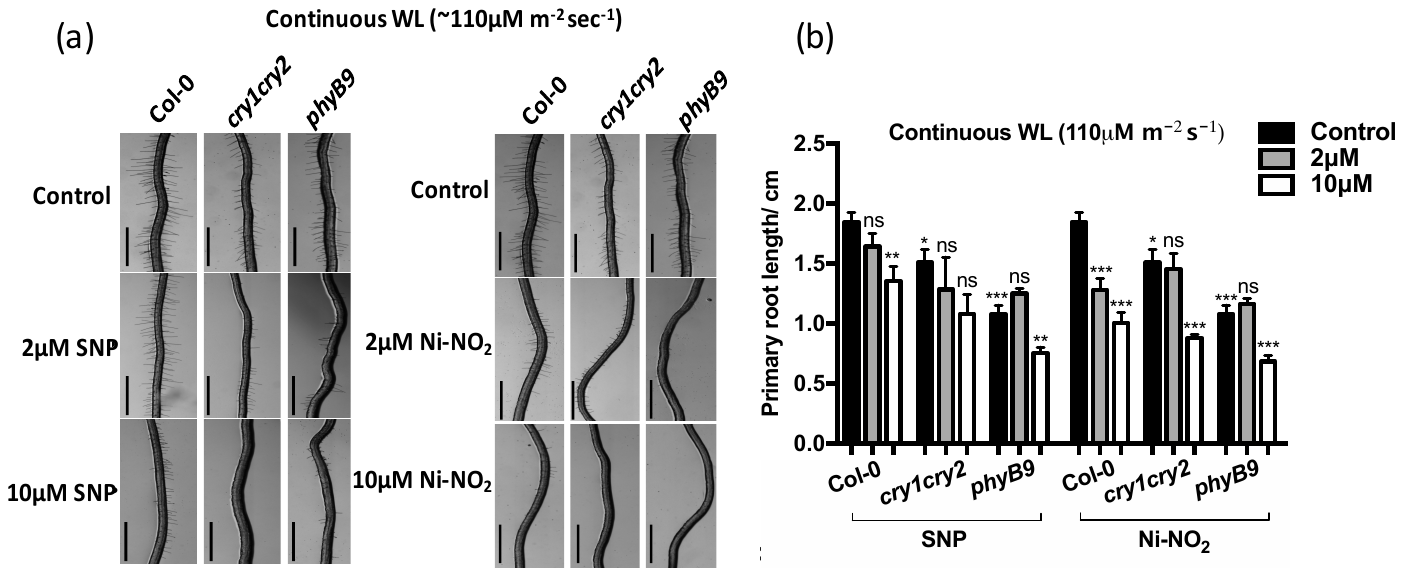

**Figure S14**. Response of WT, *cry1cry2* and *phyB-9* mutants to NO under continuous white light (a) WT, *cry1cry2* and *phyB-9* seedlings growing on medium supplemented with SNP and Ni-NO_2_ at various concentration under continuous white light at 110 *μ*M m^-2^ s^-1^ fluence. (b) Root length of WT, *cry1cry2* and *phyB-9* seedlings treated as in (a). Mean value and SE were calculated from at least 20 seedlings. Significant difference was analyzed by Student’s t-test with Welch’s corrections compared to WT under same condition are indicated by asterisks: *P <.05, ** P <.01, *** P <.001.

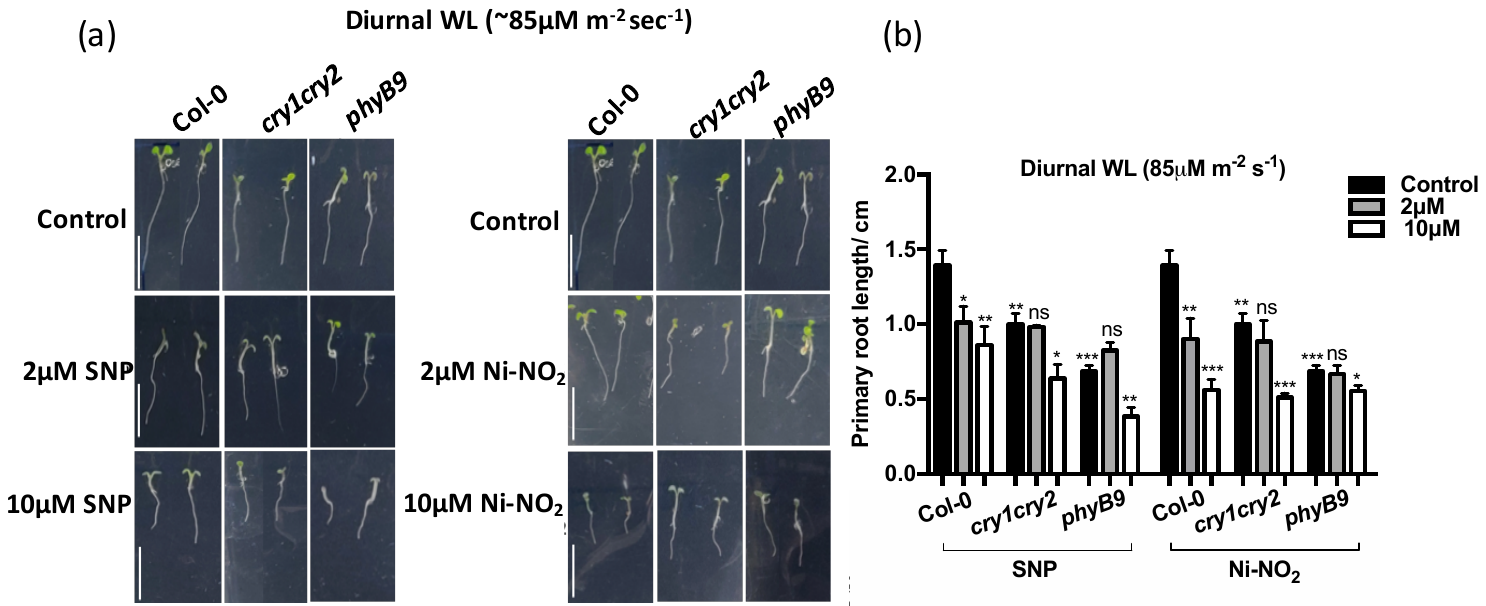

**Figure S15.** Response of WT, *cry1cry2* and *phyB-9* mutants to NO under diurnal white light (a) WT, *cry1cry2* and *phyB-9* seedlings growing on medium supplemented with SNP and Ni-NO2 at various concentration under diurnal white light at 85 *μ*M m^-2^ s^-1^ fluence. (b) Root length of WT, *cry1cry2* and *phyB-9* seedlings treated as in (a). Mean value and SE were calculated from at least 20 seedlings. Significant difference was analyzed by Student’s t-test with Welch’s corrections compared to WT under same condition are indicated by asterisks: *P <.05, ** P <.01, *** P <.001.

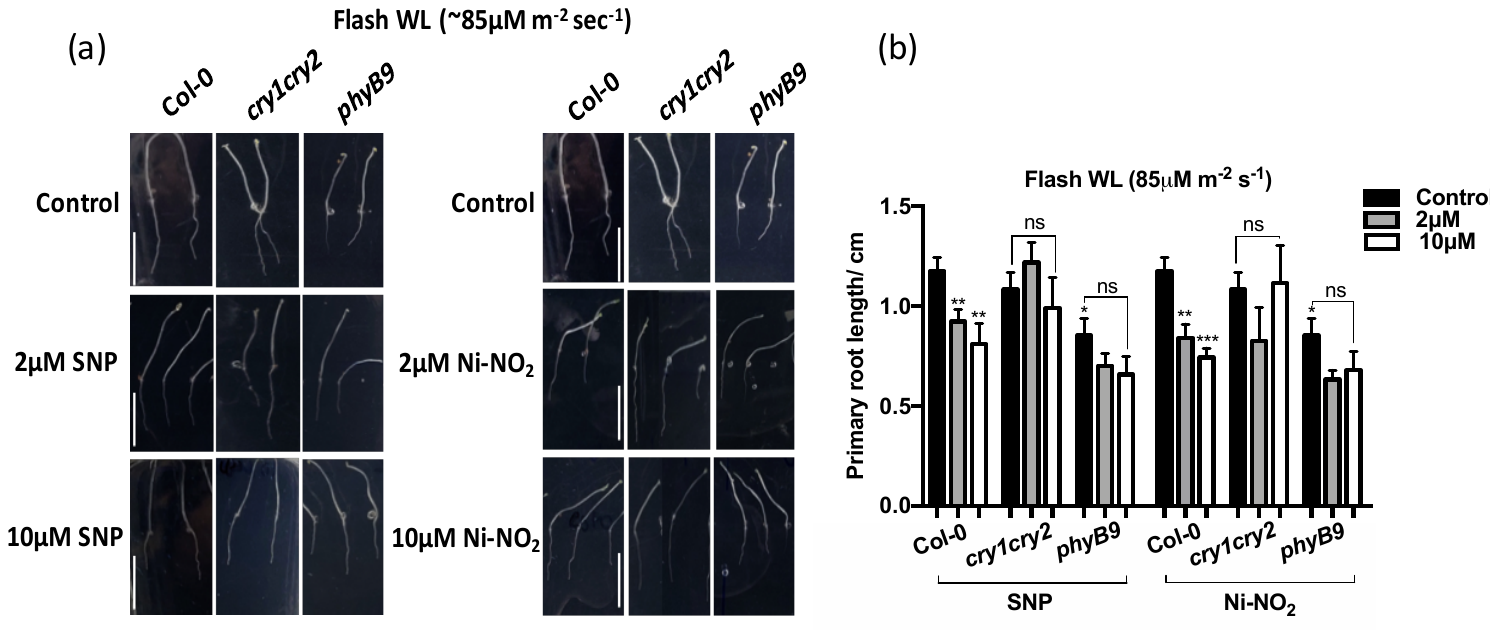

**Figure S16.** Response of WT, *cry1cry2* and *phyB-9* mutants to NO under flash white light (a) WT, *cry1cry2* and *phyB-9* seedlings growing on medium supplemented with SNP and Ni-NO_2_ at various concentration under flash white light at 85 *μ*M m^-2^ s^-1^ fluence. (b) Root length of WT, *cry1cry2* and *phyB-9* seedlings treated as in (a). Mean value and SE were calculated from at least 20 seedlings. Significant difference was analyzed by Student’s t-test with Welch’s corrections compared to WT under same condition are indicated by asterisks: *P <.05, ** P <.01, *** P <.001.

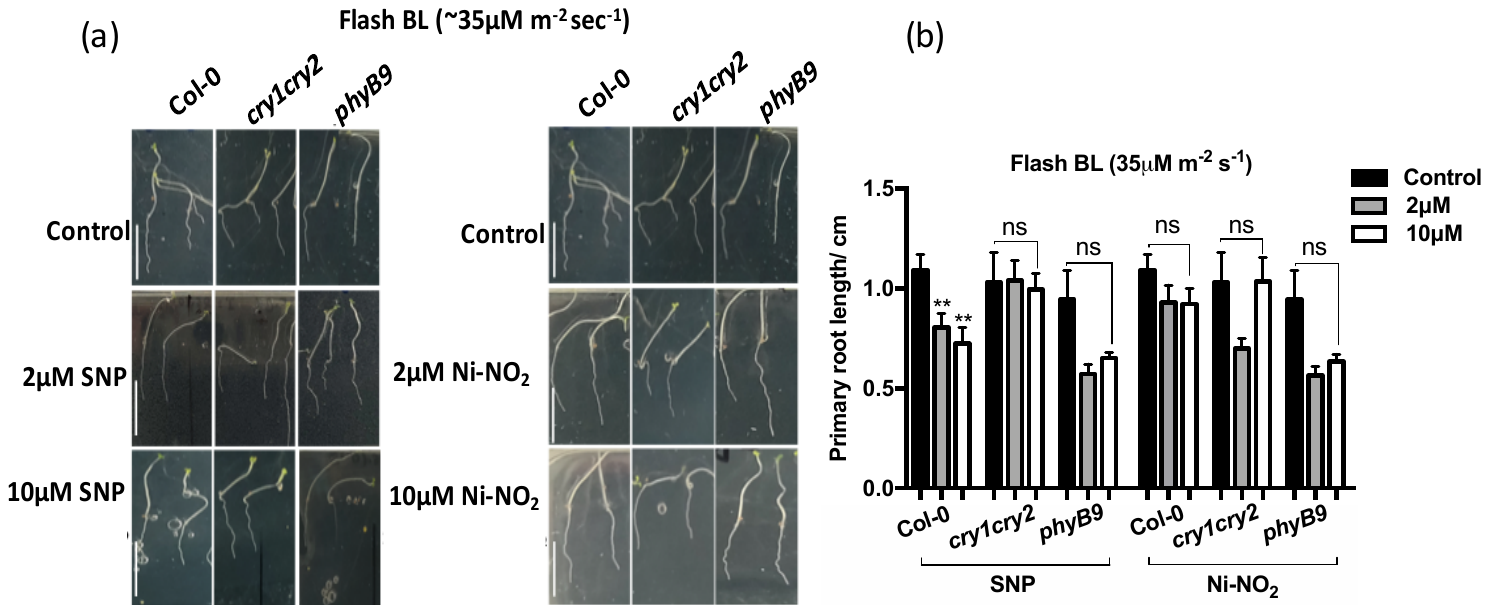

**Figure S17**. Response of WT, *cry1cry2* and *phyB-9* mutants to NO under flash blue light (a) WT, *cry1cry2* and *phyB-9* seedlings growing on medium supplemented with SNP and Ni-NO_2_ at various concentration under flash blue light at 35 *μ*M m^-2^ s^-1^ fluence. (b) Root length of WT, *cry1cry2* and *phyB-9* seedlings treated as in (a). Mean value and SE were calculated from at least 20 seedlings. Significant difference was analyzed by Student’s t-test with Welch’s corrections compared to WT under same condition are indicated by asterisks: ** P <.01.

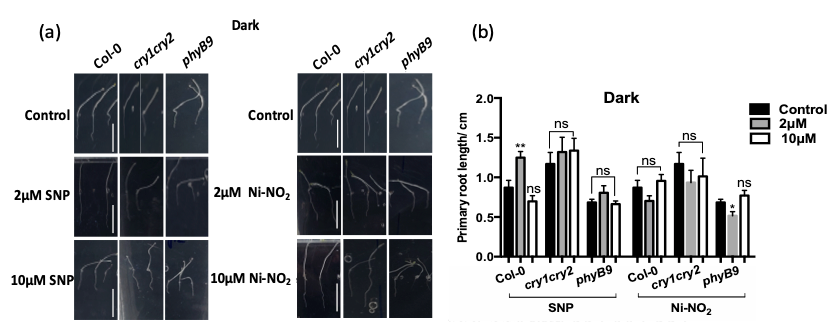

**Figure S18**. Response of WT, *cry1cry2* and *phyB-9* mutants to NO under dark (a) WT, *cry1cry2* and *phyB-9* seedlings growing on medium supplemented with SNP and Ni-NO_2_ at various concentration under dark (b) Root length of WT, *cry1cry2* and *phyB-9* seedlings treated as in (a). Mean value and SE were calculated from at least 20 seedlings. Significant difference was analyzed by Student’s t-test with Welch’s corrections compared to WT under same condition are indicated by asterisks: *P <.05, ** P <.01.

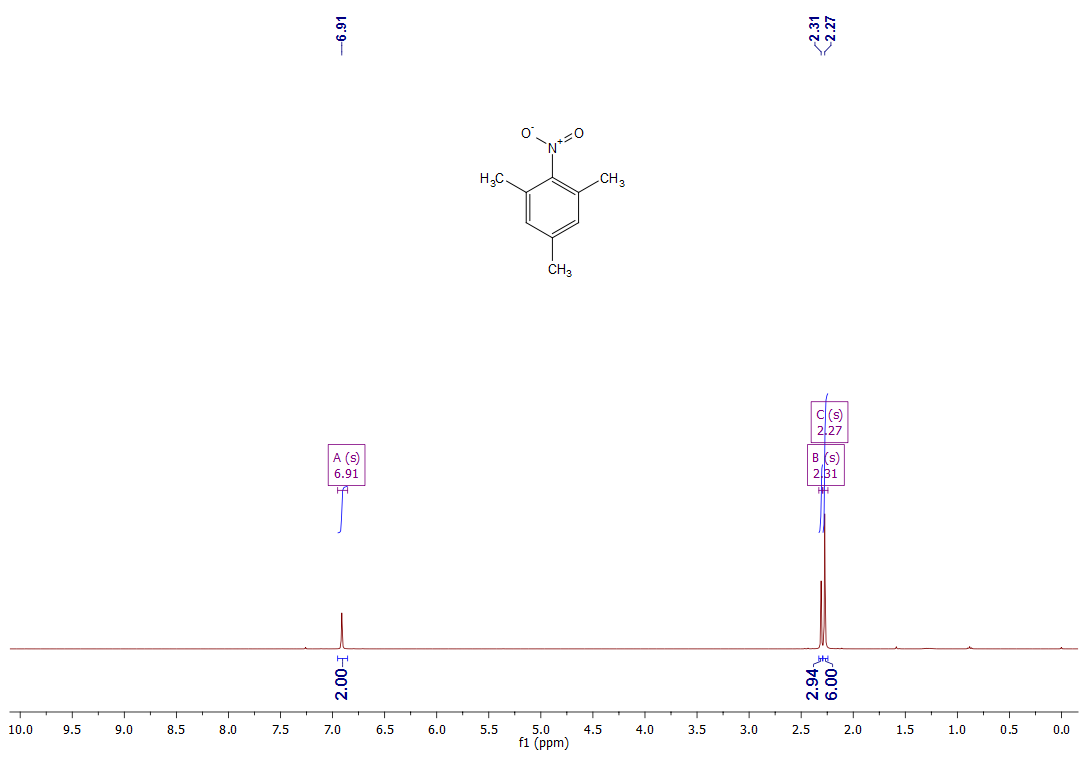

**Figure S19**. ^1^H NMR spectrum of compound **1**

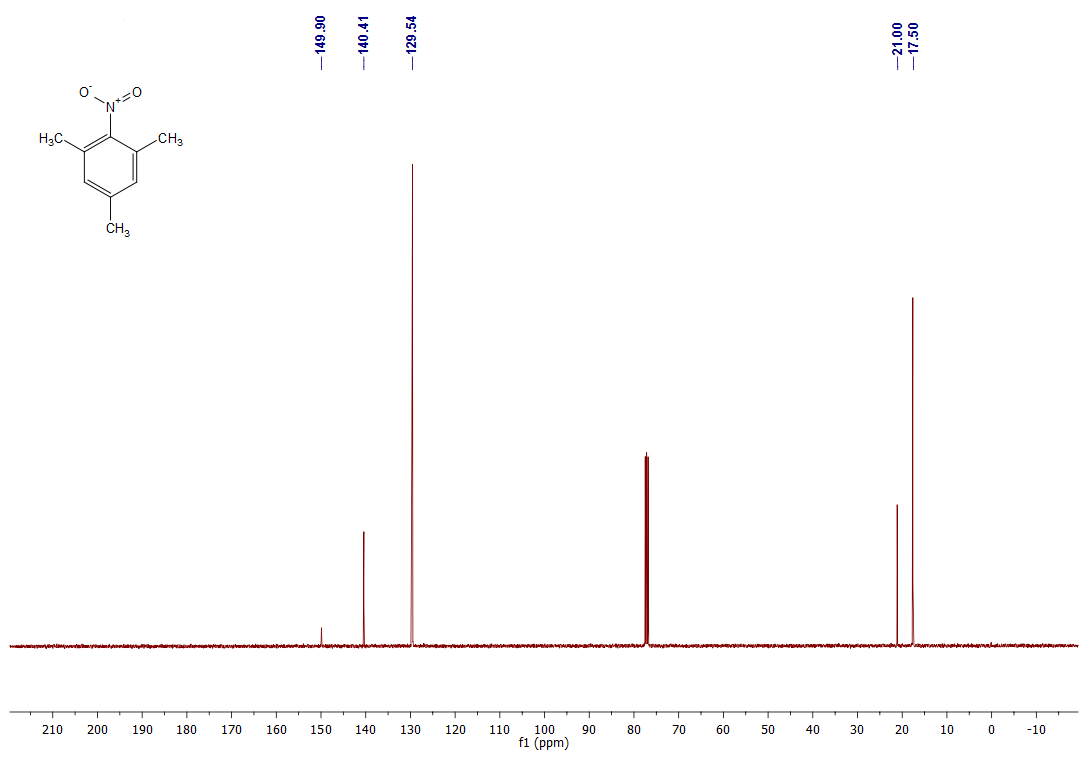

**Figure S20**. ^13^C NMR spectrum of compound **1**

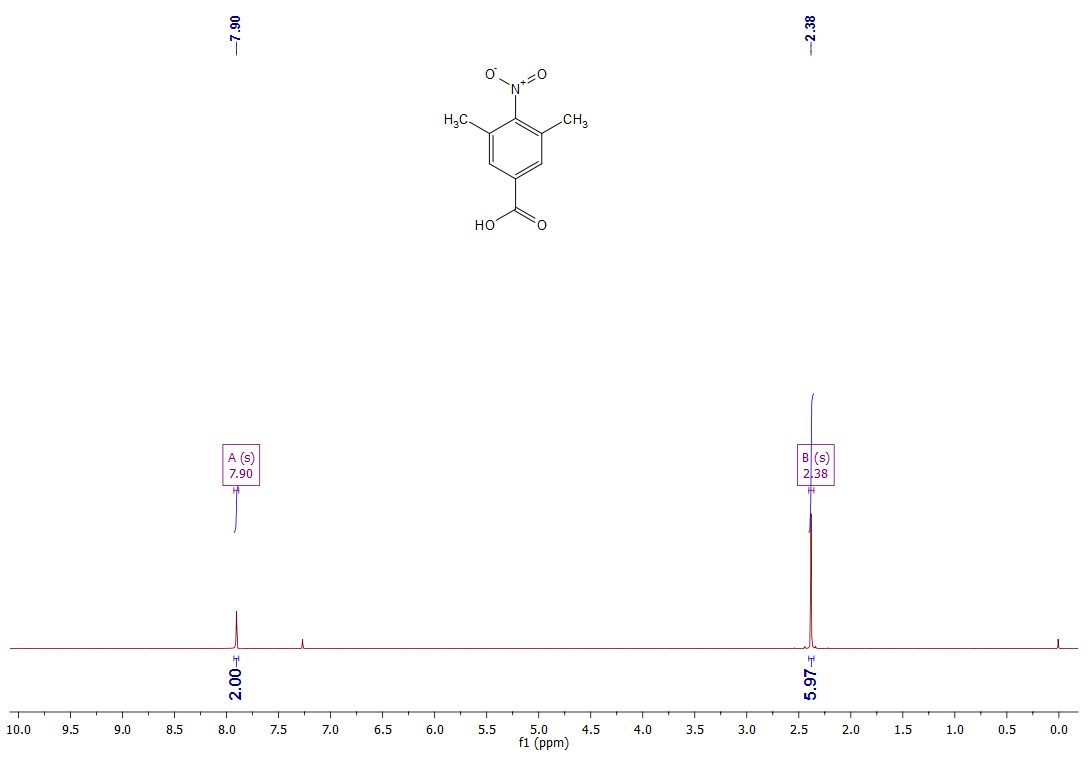

**Figure S21**. ^1^H NMR spectrum of compound **2**

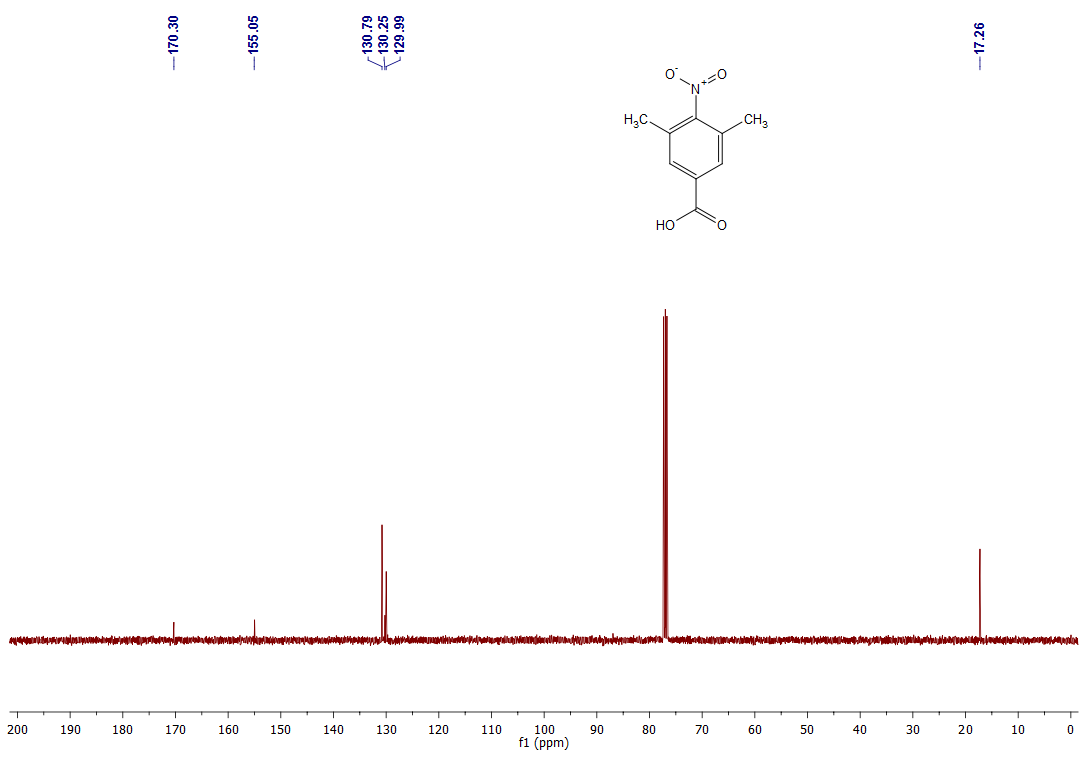

**Figure S22**. ^13^C NMR spectrum of compound **2**

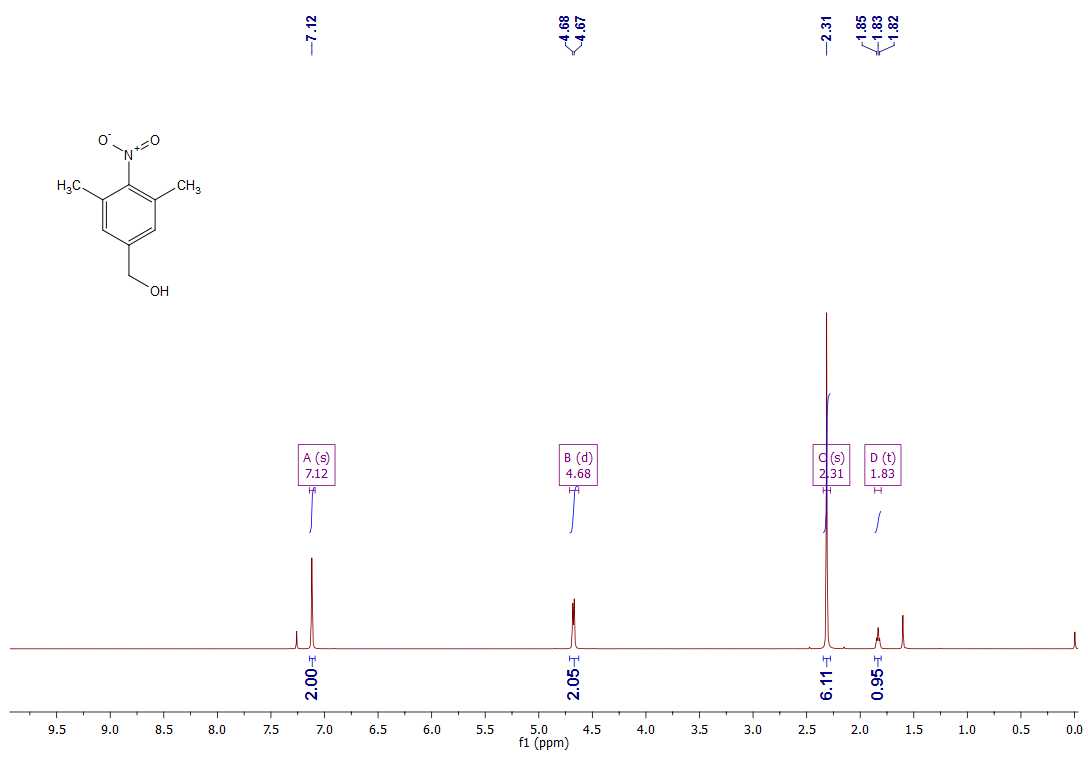

**Figure S23**. ^1^H NMR spectrum of compound **3a**

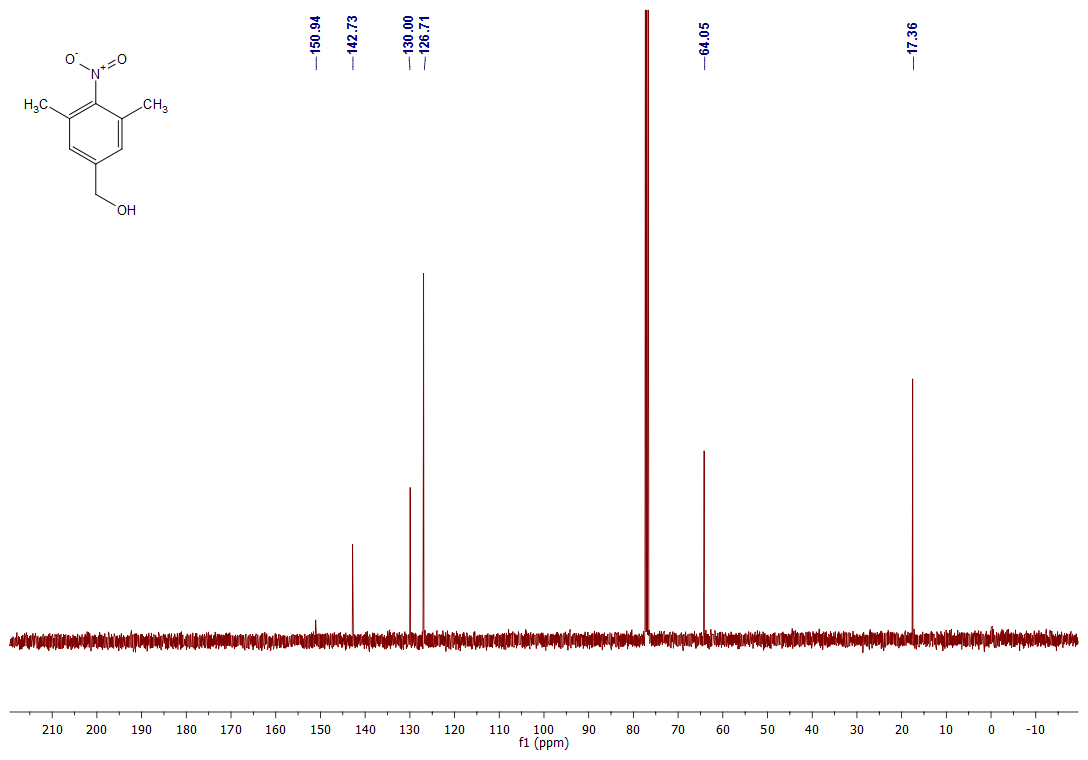

**Figure S24**. ^13^C NMR spectrum of compound **3a**

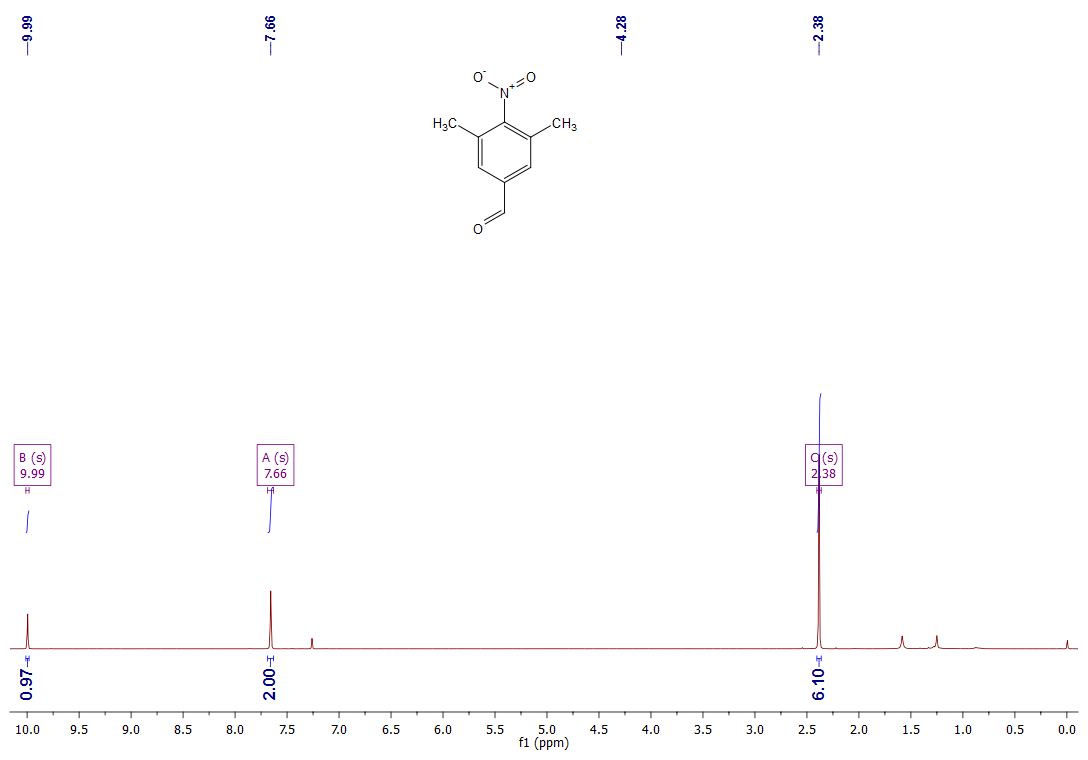

**Figure S25**. ^1^H NMR spectrum of compound **3b**

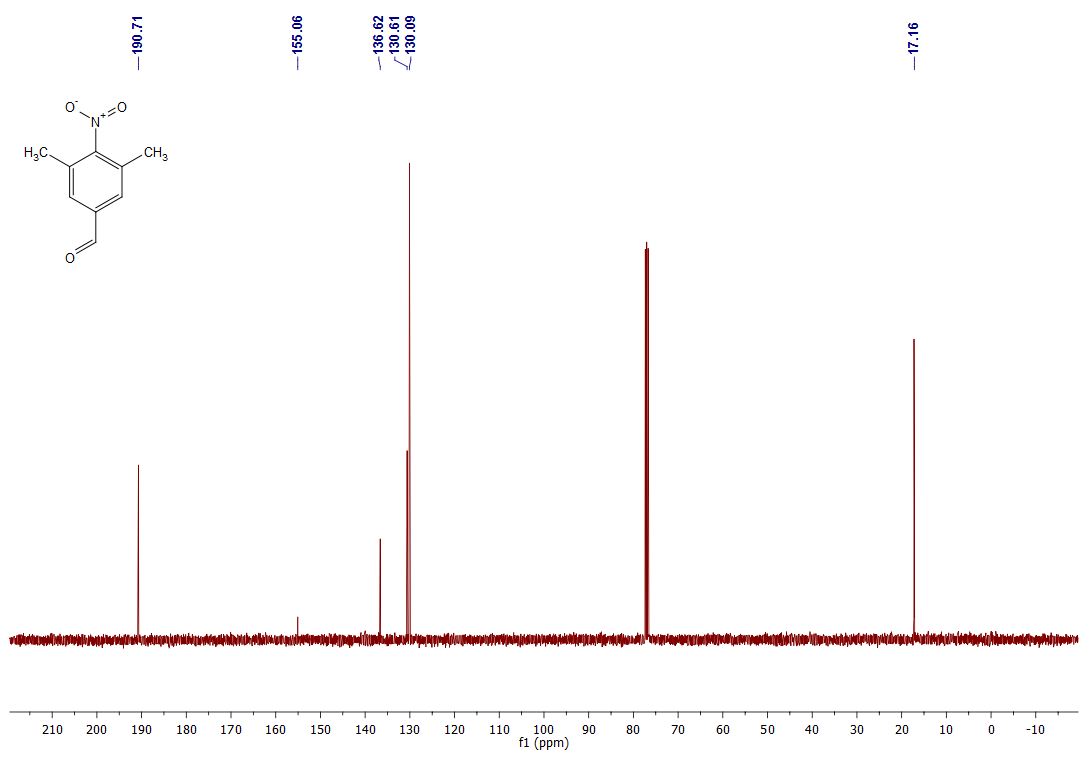

**Figure S26**. ^13^C NMR spectrum of compound **3b**

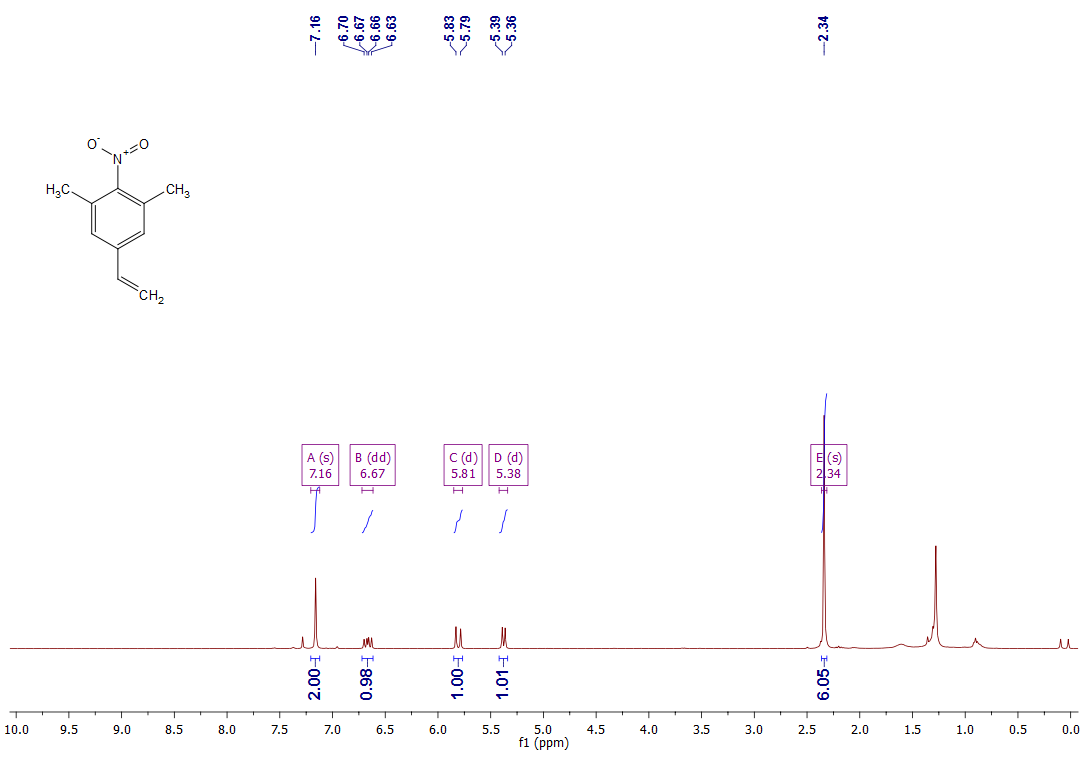

**Figure S27**. ^1^H NMR spectrum of compound **4**

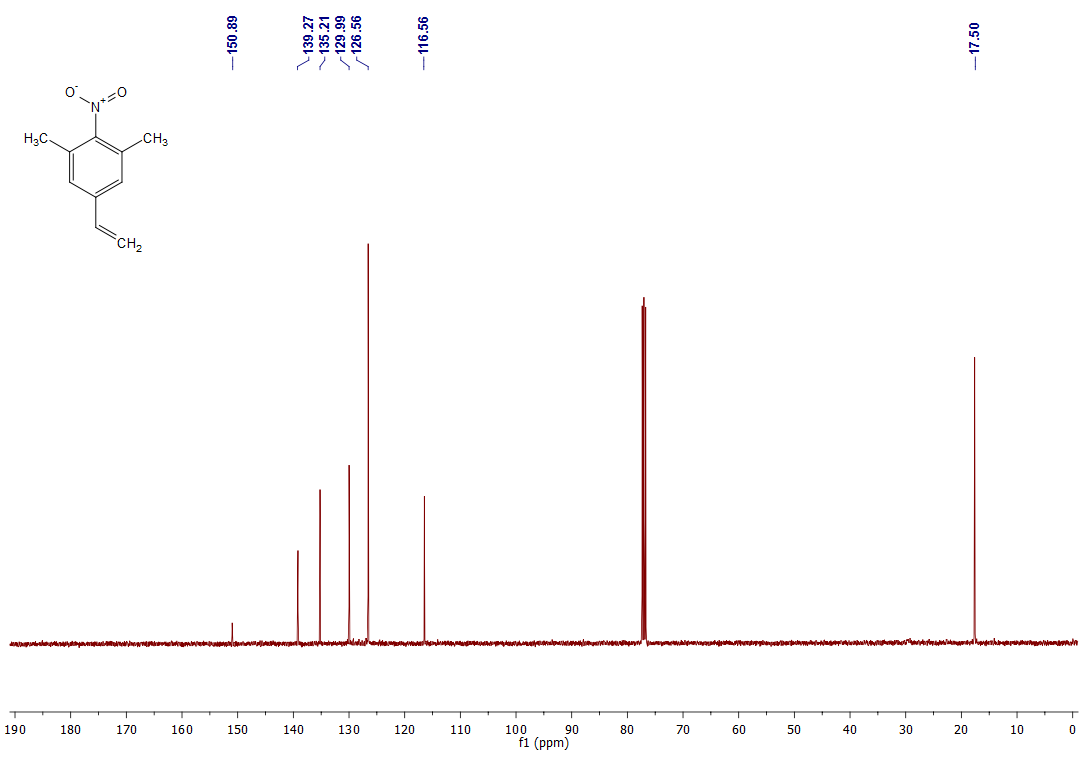

**Figure S28**. ^13^C NMR spectrum of compound **4**

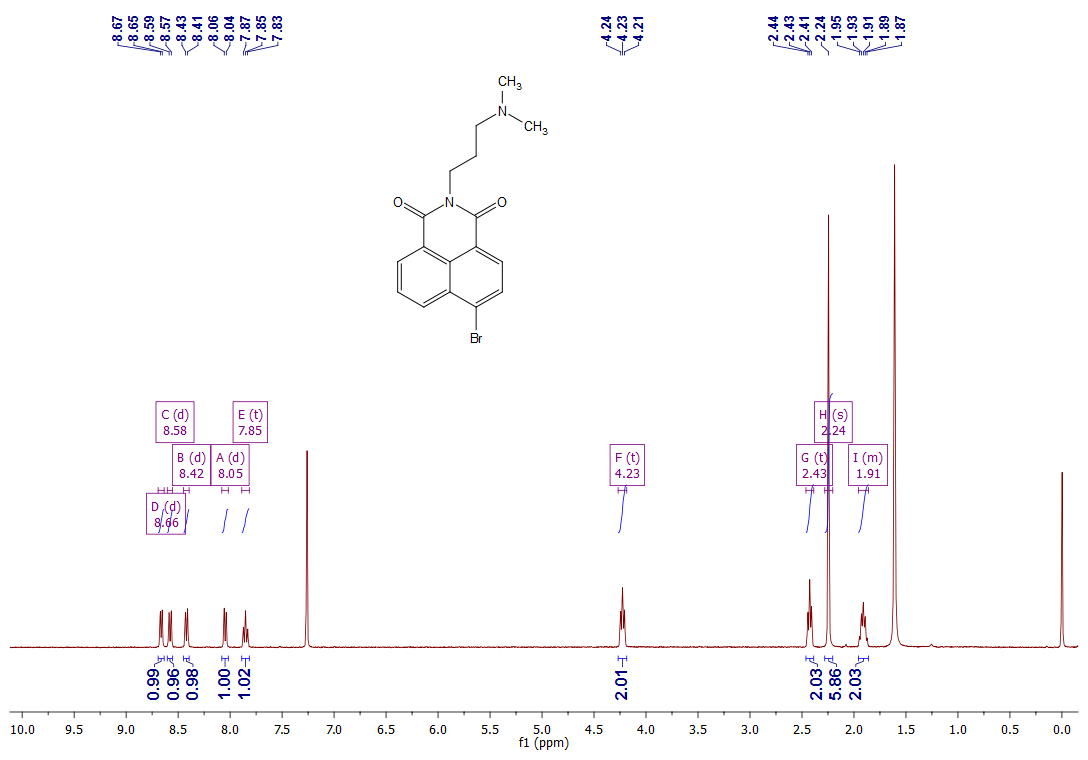

**Figure S29**. ^1^H NMR spectrum of compound **5**

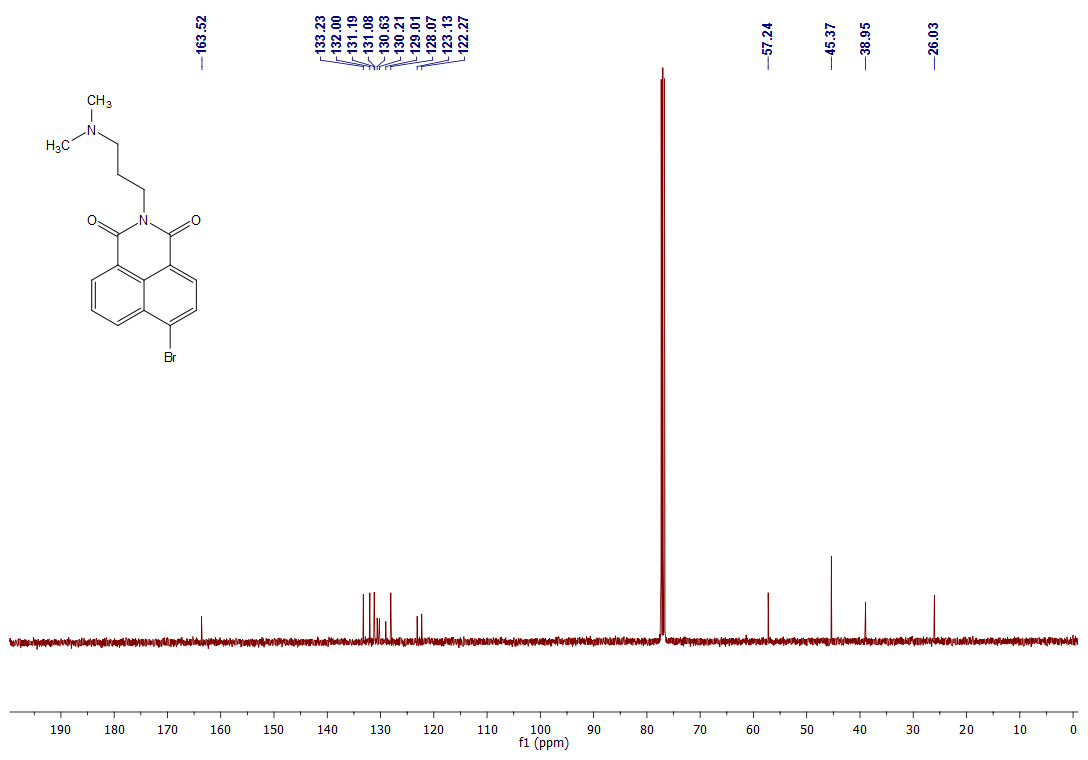

**Figure S30**. ^13^C NMR spectrum of compound **5**

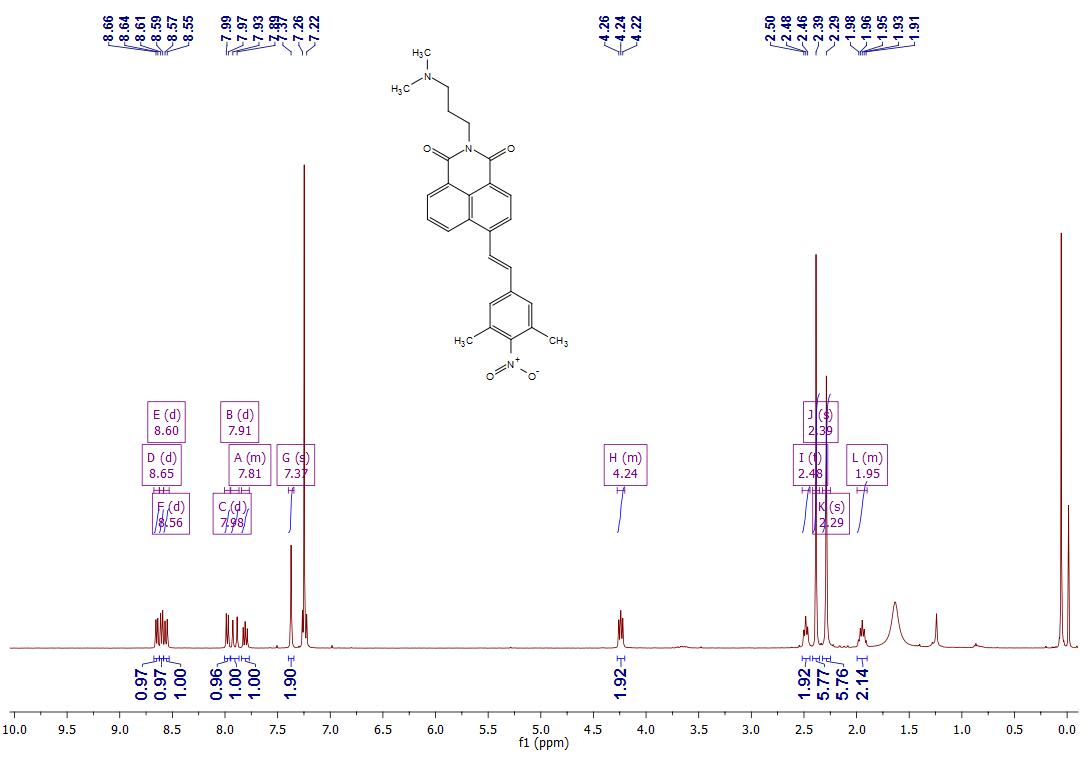

**Figure S31**. ^1^H NMR spectrum of compound **Ni-NO_2_**

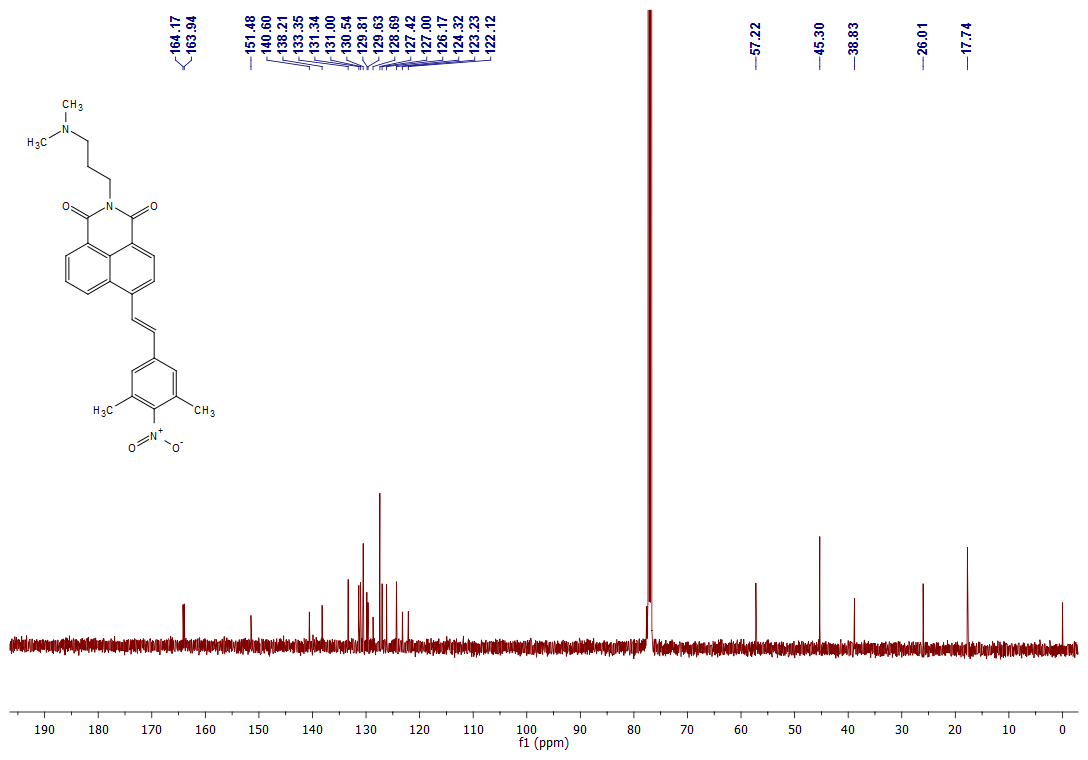

**Figure S32**. ^13^C NMR spectrum of compound **Ni-NO_2_**

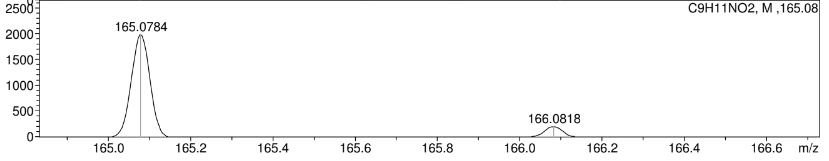

**Figure S33**. APCI HRMS mass spectrum of compound **1**

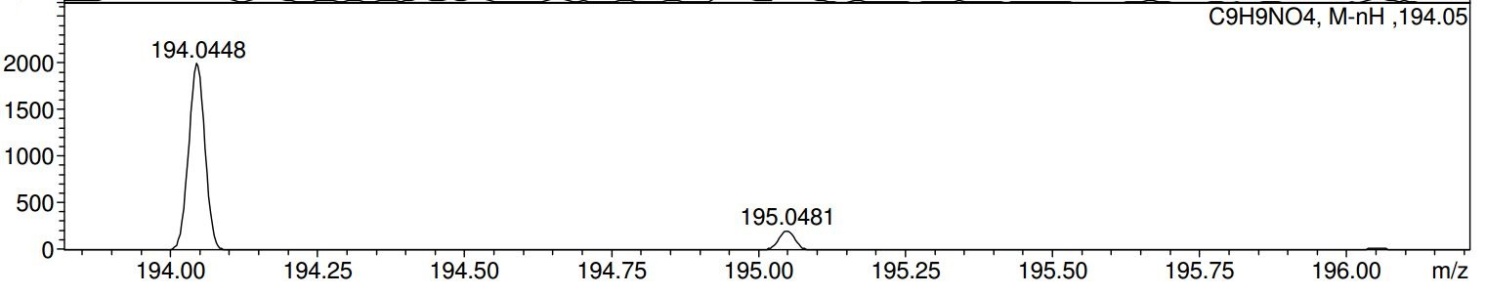

**Figure S34**. ESI-mass spectrum of compound **2**

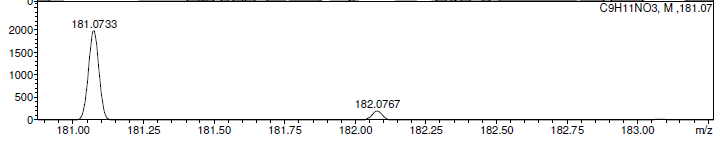

**Figure S35**. APCI HRMS mass spectrum of compound **3a**

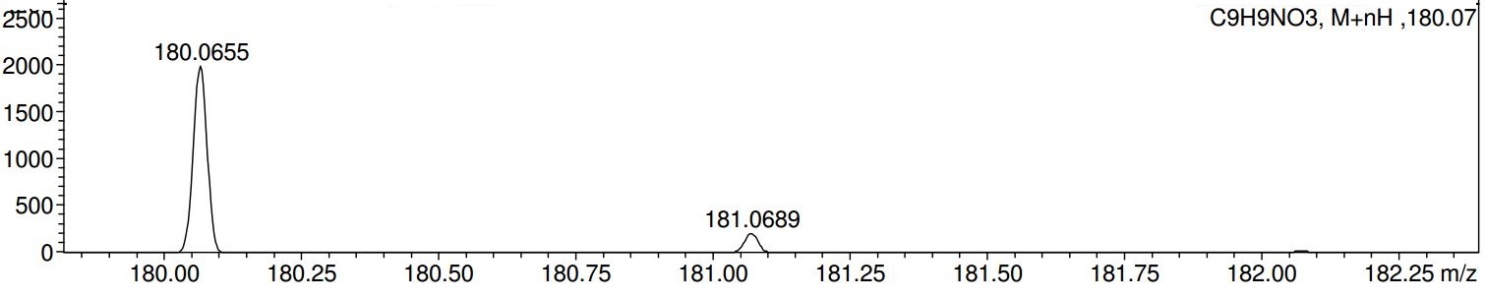

**Figure S36**. APCI HRMS mass spectrum of compound **3b**

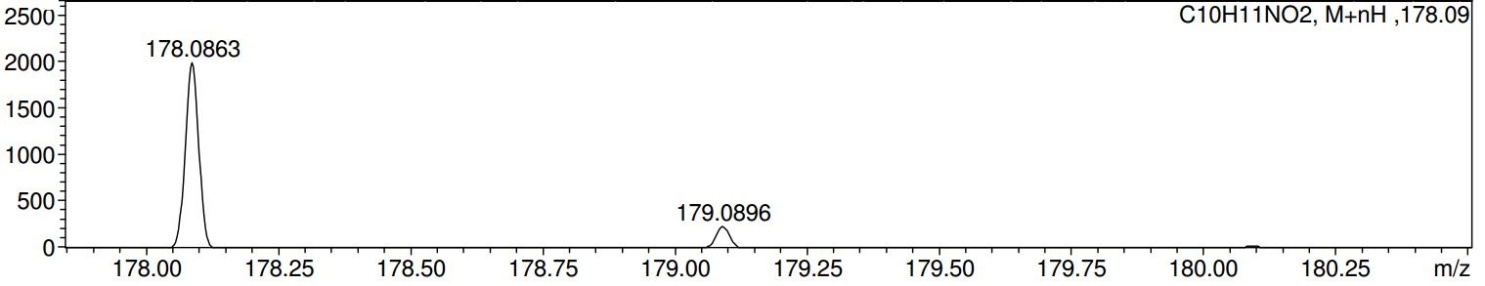

**Figure S37**. APCI HRMS mass spectrum of compound **4**

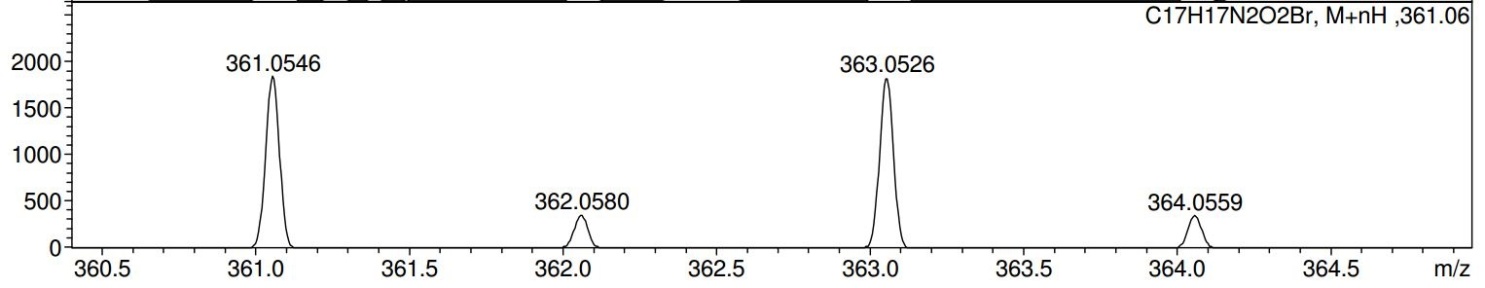

**Figure S38**. APCI HRMS mass spectrum of compound **5**

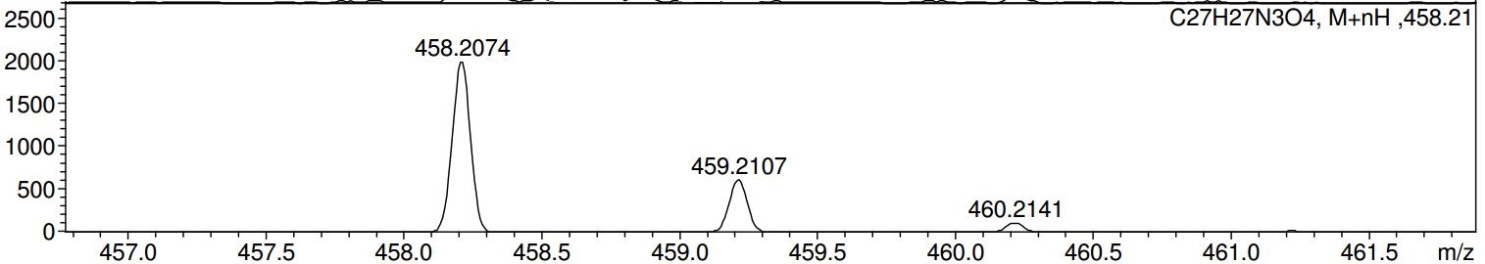

**Figure S39**. APCI HRMS mass spectrum of **Ni-NO_2_**
